# Post-replicative initial expression of the cell fate regulator *PAX6* during neuroectoderm differentiation

**DOI:** 10.1101/2025.05.30.657125

**Authors:** Song Hu, Rongao Kou, Zhuojie Su, Guanchen Li, Shutao Qi, Yanxiao Zhang, Haifeng Wang, Ling-Ling Chen, Hongtao Yu

## Abstract

The development of multicellular organisms requires precise coordination between cell division and differentiation. Cell division generates the necessary number of cells, while differentiation creates distinct cell identities, forming tissues and organs. The transcription factors SOX2 and PAX6 specify neuroepithelial cells, the earliest neural progenitor cells (NPCs) during brain development. How lineage specification is coordinated with the cell cycle is not fully understood. Here, we show that *PAX6* expression occurs during a narrow time window––between 48 and 72 hours––after neural induction of human embryonic stem cells (ESCs). Flow cytometry analyses and time-lapse imaging further show that *PAX6* expression starts during the G2 phase of the cell cycle. We identify a novel 500-bp *PAX6* promoter that drives its G2-specific expression. PAX6 expression is independent of known regulators of cell-cycle-dependent transcription, suggesting the existence of a novel mechanism. S-phase block by hydroxyurea prevents PAX6 expression and differentiation into NPC. Thus, NPC fate specification is coupled to cell cycle progression and occurs after the completion of DNA replication. This post-replicative lineage commitment ensures the creation of two daughter cells of identical cell fate following cell division.

## Introduction

The development of multicellular organisms requires precisely timed and spatially coordinated processes that balance cell division and differentiation. Cell division generates enough cells necessary for building tissues and organs, while differentiation assigns distinct identities to these cells, enabling functional specialization^1^. This balance is dynamically regulated by extracellular signals, such as WNT, BMP, SHH, and FGF families of proteins, which form morphogen gradients to instruct cells based on their spatial positioning^2^. These signals activate intracellular pathways that regulate transcription factor networks, which in turn govern cell identity^3^.

For differentiation into terminally differentiated, functional cell types that no longer divide, such as neurons, the precursor cells often undergo cell cycle exit and activate the expression of cell-fate-determining genes to commit to a particular cell type^4,5^. During early embryogenesis, however, pluripotent stems cells (PSCs) differentiate into multipotent precursors of specialized lineages that can themselves under active cell divisions^6,7^. During this situation, it is unclear whether cell fate decisions are coupled to cell cycle progression, and if so, during which cell cycle phase the lineage commitment is made.

The mammalian nervous system develops from neuroepithelial cells, the earliest neural progenitor cells (NPCs) lining the neural tube^8^. These cells exhibit apical-basal polarity and undergo symmetric divisions to expand the progenitor pool before giving rise to radial glia that can both self-renew and undergo asymmetric divisions to produce neurons^8,9^. The transcription factors SOX2 and PAX6 regulate neuroectodermal specification^10–12^. Mutations in SOX2 and PAX6 cause severe eye and brain malformations^11–13^.

The cell cycle is tightly regulated by phase-specific transcriptional programs^14^. Cell cycle-dependent transcription is mediated by conserved regulatory elements in gene promoters, which recruit transcription factors in a temporally controlled manner. The DREAM complex (DP, RB-like, E2F, and MuvB) is a key transcriptional silencer of activation of mitotic genes. During cell cycle progression, the DREAM complex is disrupted, leading to the formation of the related MMB-FOXM1 complex that contains B-MYB and FOXM1^14^. The MMB-FOXM1 complex binds to promoter regions of genes encoding mitotic regulators, such as cyclin B, PLK1, and Aurora kinases, through interactions with cell cycle homology region (CHR) elements or CCAAT-box motifs^15–17^. At the G2–M transition, CDK1 and PLK1 phosphorylate and activate the MMB-FOXM1 complex, promoting the transcription of mitotic genes^14,18^.

Emerging evidence suggests that the cell cycle is not merely a passive timer but an active regulator of differentiation. In mammalian systems, PSCs lengthen their cell cycle, particularly the G1 phase, during differentiation^19^. G1 cyclins, cyclin D1–3, alter the propensity of PSCs to differentiate into different lineages through regulating the TGFβ pathway^20^. It has been proposed that cell-cycle-dependent mechanisms restrict the activation of developmental genes in PSCs to the G1 phase^21^.

Previous work has established an *in vitro* differentiation protocol to differentiate PSCs into NPCs by inhibiting TGFβ and BMP pathways^22^. In this study, using this dual SMAD inhibition (dSMADi) differentiation protocol, we differentiated human embryonic stem cells (ESCs) into NPCs and monitored the expression of the NPC marker PAX6 during the process using flow cytometry and live-cell imaging of cells expressing FUCCI (Fluorescent Ubiquitination-based Cell Cycle Indicator)^20,23^. PAX6 activation is confined to a narrow temporal window––48 to 72 hours after differentiation initiation. Its initial expression coincides with the G2 phase of the cell cycle. Intriguingly, a 500-base-pair (bp) region of the *PAX6* promoter drives its G2-specific expression in a mechanism that does not appear to involve the DREAM complex. S-phase arrest by the DNA replication inhibitor hydroxyurea abolishes PAX6 expression and blocks NPC specification. These findings reveal a novel mechanism where NPC commitment is contingent on cell cycle progression, with lineage determination occurring post-replication to ensure the generation of two NPCs upon division.

## Results

### Neuroectoderm cell fate transition occurs in a short time window during *in vitro* differentiation

To investigate the cell fate transition of neuroectoderm *in vitro*, we differentiated H9 ESCs into NPCs using the established dSMADi neural induction protocol^22^, which required 18 days to differentiate ESCs into NPCs. To assess the robustness of this neural induction system, we collected cells at various timepoints during differentiation and subjected them to Western blotting (Extended Data Fig. 1a). As SOX2 was expressed highly in both ESCs and NPCs^10,24^, the established NPC marker PAX6 was used to track NPC identity. The PAX6 protein was absent in ESCs and accumulated during neural induction. We then characterized the transcriptional profiles of differentiated NPCs using bulk RNA sequencing (RNA-seq). Pathway analysis revealed that, compared to ESCs, upregulated differentially expressed genes (DEGs) in NPCs at D18 were predominantly associated with central nervous system development (Extended Data Fig. 1b), confirming successful NPC generation. To evaluate their differentiation potential, we differentiated NPCs into forebrain neurons. Immunofluorescence (IF) demonstrated widespread expression of the neuronal marker TUJ1 in NPC-derived cells post-differentiation (Extended Data Fig. 1c). RNA-seq showed that upregulated DEGs in differentiated cells were linked to synaptic formation and neurogenesis (Extended Data Fig. 1d), validating functional neuronal differentiation. Collectively, these results confirm the robust transition from ESCs to NPCs in our *in vitro* system, enabling subsequent mechanistic studies of cell fate transitions.

Next, we sought to pinpoint the initial timing of NPC fate specification. Samples were collected daily during the first 6 days of neural induction and analyzed for PAX6 expression with Western blotting, IF, and FACS. PAX6 protein levels rose sharply between D2 and D3 (Fig. 1a), corroborated by IF and FACS quantification showing PAX6^+^ cells increasing from ∼5% at D2 to ∼50% by D3 (Fig. 1b-d). To confirm whether this time window indeed corresponded to the ESC-to-NPC transition, we also examined pluripotency markers (NANOG and OCT4) in addition to PAX6 by IF. NANOG disappeared by D2, while OCT4 persisted at low levels on D2 but was absent by D3 (Fig.1e). PAX6 expression initiated on D2 and further increased by D3. Thus, D2 represents the intermediate phase of NPC fate specification.

**Fig. 1.**
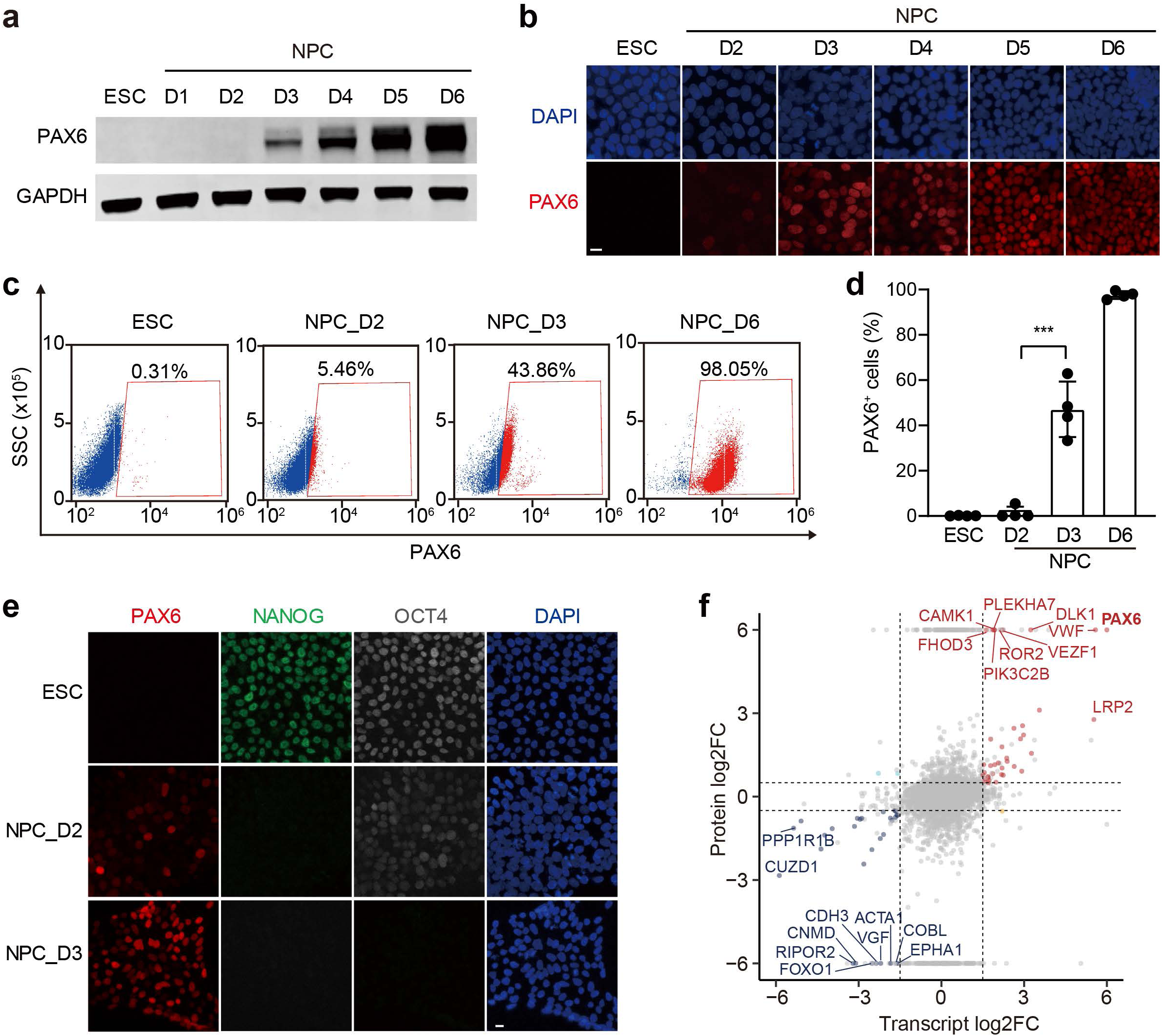
| The ESC–NPC cell fate transition occurs between day 2 (D2) and D3 of neural induction. **a**, Western blot analysis of PAX6 protein levels in ESCs and NPCs during D1 to D6 of neural induction. GAPDH was used as the loading control. **b**, Images of ESCs and NPCs at indicated times of neural induction stained with anti-PAX6 antibody and DAPI. Scale bar, 10 μm. **c**, FACS analysis of the percentage of PAX6^+^ cells in ESCs and NPCs at indicated times of neural induction. **d**, Quantification of the percentage of PAX6^+^ cells in **c**. n=4 independent experiments. **e**, Images of ESCs and NPCs at D2 and D3 of neural induction stained with DAPI and antibodies against NANOG, OCT4, and PAX6. Scale bar, 10 μm. **f**, Scatter plot showing the correlation of transcriptomics and proteomics data of differentially expressed genes and proteins between NPC_D3 and NPC_D2. The horizontal and vertical dotted lines indicate significance boundaries (transcriptomics: p value(0.01 and |log2FC|>1.5; proteomics: p value(0.05 and |log2FC|>0.5).

To probe gene expression dynamics during this transition, we used bulk RNA-seq and mass spectrometry to profile transcriptomic and proteomic changes. RNA-seq revealed extensive transcriptional remodeling between D2 and D3 (1,037 upregulated and 1,834 downregulated DEGs; Extended Data Fig. 2a), with upregulated genes enriched in neurogenesis and brain development pathways (Extended Data Fig. 2b). Proteomic analysis identified fewer differentially expressed proteins (247 upregulated and 302 downregulated proteins; Extended Data Fig. 2c). Upregulated proteins similarly mapped to neurodevelopmental pathways (Extended Data Fig. 2d). The transcript-protein concordance was partial (Extended Data Fig. 2e,f). Notably, *PAX6* expression exhibited concurrent upregulation patterns in both transcriptomic and proteomic datasets (Fig. 1f), suggesting that its transcriptional activation might, to some degree, account for PAX6 expression after D2 of neural induction.

Time-series proteomic clustering revealed four distinct protein expression patterns during ESC-to-NPC differentiation (Extended Data Fig. 2g). Clusters 2 and 3, representing upregulated proteins, were enriched in neural development pathways, again consistent with neural induction (Extended Data Fig. 2h). The four clusters were also enriched for additional pathways, including ion transport, RNA processing, and cell cycle regulation. These proteomic changes highlight the complexity of cell fate transitions and the involvement of multiple cellular processes, including the cell cycle.

### PAX6 expression initiates in G2 during NPC cell-fate transition

We next analyzed PAX6 expression from D2 to D3 during neural induction. Western blotting revealed a gradual increase in PAX6 protein levels during differentiation from 48 to 72 hours (Fig. 2a and Extended Data Fig. 3a). Quantitative PCR demonstrated a similar upregulation of *PAX6* mRNA levels (Extended Data Fig. 3b). FACS analysis further confirmed a continuous increase in the proportion of PAX6^+^ cells over this period (Fig. 2b and Extended Data Fig. 3c). Notably, the earliest PAX6^+^ cells detected by FACS had 4N DNA content, indicating that they were in the G2/M phase (Fig. 2b). To further pinpoint whether PAX6 expression initiated in G2 or mitosis, we employed the FUCCI system to simultaneously monitor PAX6 expression and the cell cycle status from 46 to 72 hours using IF (Fig. 2c and Extended Data Fig. 3d). PAX6^+^ cells first emerged at 46 hours of differentiation, with their percentages gradually increasing. Strikingly, nearly all early PAX6^+^ cells were GEMININ^+^ and CDT1^−^ with intact nuclei, indicating that these cells were in G2 (Fig. 2c,d).

**Fig. 2.**
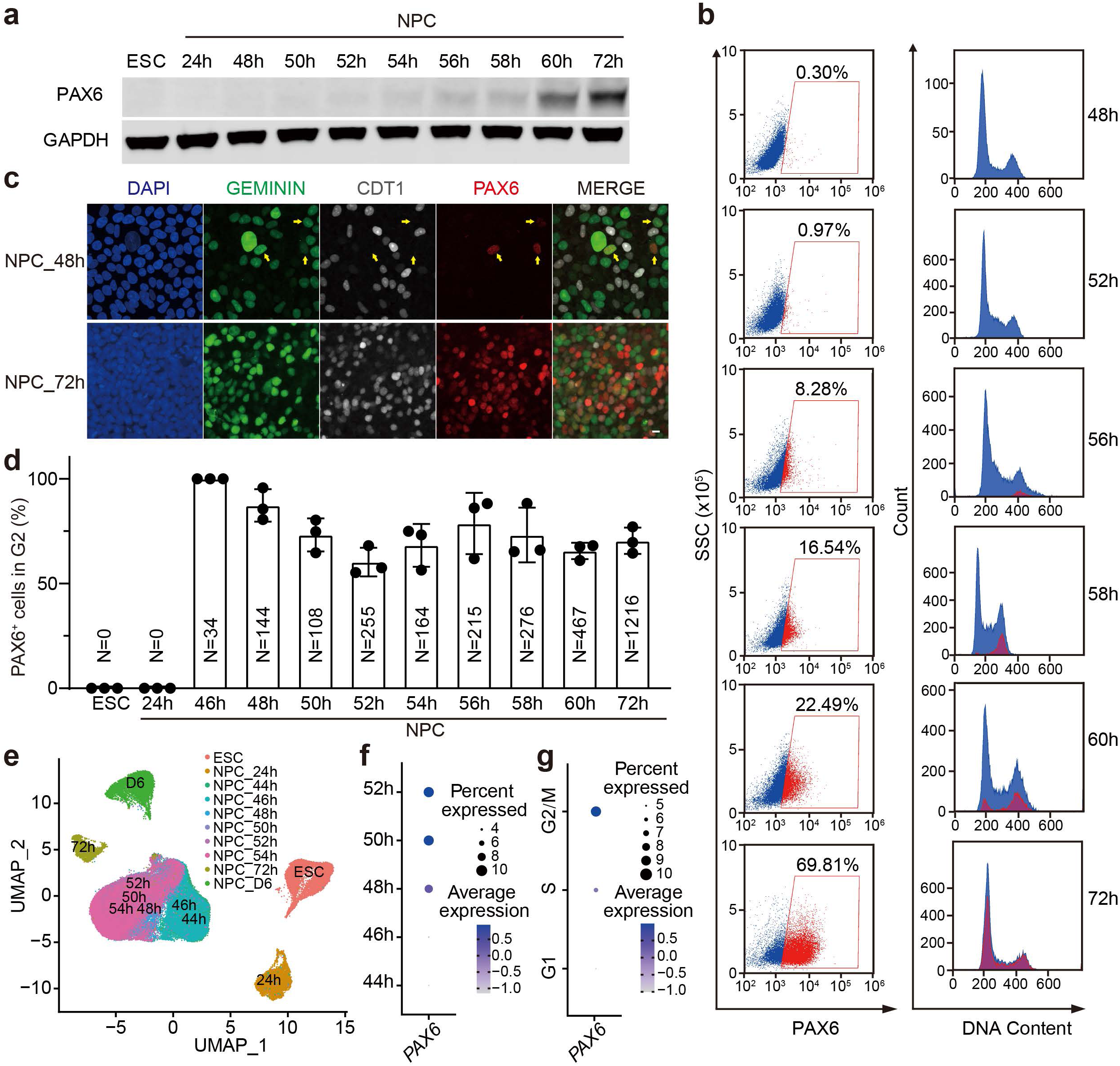
| Initial PAX6 expression occurs at G2 phase during ESC–NPC cell fate transition. **a**, Western blot analysis of PAX6 protein levels in ESCs and NPCs at different timepoints (24–72 hours) of neural induction. GAPDH was used as the loading control. **b**, FACS analyses of the percentage of PAX6^+^ cells (left panels) and the cell cycle status (right panels) in NPCs at different timepoints (48–72 hours) during neural induction. **c**, Images of FUCCI NPCs at 48 and 72 hours of neural induction stained with DAPI and the PAX6 antibody (red). The cell cycle status was determined by the FUCCI reporters (GEMININ, green; CDT1, white). Scale bar, 10 μm. **d**, Quantification of the percentage of PAX6^+^ cells at the G2 phase in **c**. Data are presented as mean ± SD, n = 3 independent experiments. N indicates the total number of PAX6^+^ cells counted in each group from the 3 independent experiments. **e**, UMAP plot of single-cell RNA sequencing (scRNA-seq) data of ESCs and NPCs at different times of neural induction. **f**, Dot plot showing *PAX6* expression at different timepoints (44–52 hours) of neural induction based on the scRNA-seq data in **e**. **g**, Dot plot showing the cell cycle distribution of all *PAX6*^+^ cells during 24–52 hours of neural induction based on integrated scRNA-seq data.

To determine whether *PAX6* mRNA also emerged during G2, we performed single-cell RNA sequencing (scRNA-seq) to profile transcriptional dynamics during ESC-to-NPC differentiation. UMAP clustering revealed distinct distributions of ESC, NPC_24h, NPC_72h, and NPC_D6 populations, whereas NPC_46h to NPC_54h samples clustered together and exhibited overlapping profiles (Fig. 2e), consistent with the gradual progression of neural induction during that time window. Analysis of marker genes showed rapid downregulation of ESC markers *NANOG* and *OCT4* and progressive *PAX6* upregulation upon neural induction (Extended Data Fig. 4a). *SOX2*, expressed in both ESCs and NPCs, was markedly elevated in NPCs, consistent with prior studies^10,25^. The percentage of *PAX6*-expressing cells gradually increased between 48–52 hours of differentiation (Fig. 2f). Cell cycle analysis of these *PAX6*-expressing cells revealed that they were predominantly in G2 (Fig. 2g and Extended Data Fig. 4b). Thus, initial *PAX6* expression at the RNA level also occurs during G2.

In addition to *PAX6*, other neural development genes exhibited cell cycle-specific expression patterns during neural induction. For example, *MAFK* and *FOXA2* were mainly expressed in G1 (Extended Data Fig. 4c), whereas *TEAD2* and *ZEB1* were expressed in G2 (Extended Data Fig. 4d). These findings suggest that G2-initiated transcription is not unique to *PAX6,* but represents a broader regulatory mechanism for multiple genes during cell fate transitions.

### A novel 500-bp promoter drives G2-specific PAX6 expression

*PAX6* is known to use three promoters (P0, P1, and Pα) for expression, with P1 implicated in early neural development^26,27^. For a silenced gene to become active, chromatin remodeling must increase the accessibility of its promoter to transcription factor binding^28^. We performed ATAC-seq to map chromatin accessibility changes during neural induction (Fig. 3a)^29^. The chromatin accessibility of promoters of *OCT4* and *NANOG* markedly decreased by D1 of neural induction, consistent with their role in maintaining pluripotency (Extended Data Fig. 5a). Genome-wide analysis identified numerous chromatin regions transitioning from closed to open states during neural induction (Extended Data Fig. 5b), with associated genes enriched for neural development pathways. Notably, the accessibility of the *PAX6* P1 promoter region markedly increased between D2 and D3 (Fig. 3a), consistent with the expression timing of *PAX6*. This finding suggested that P1 might mediate cell cycle-dependent *PAX6* transcription.

**Fig. 3.**
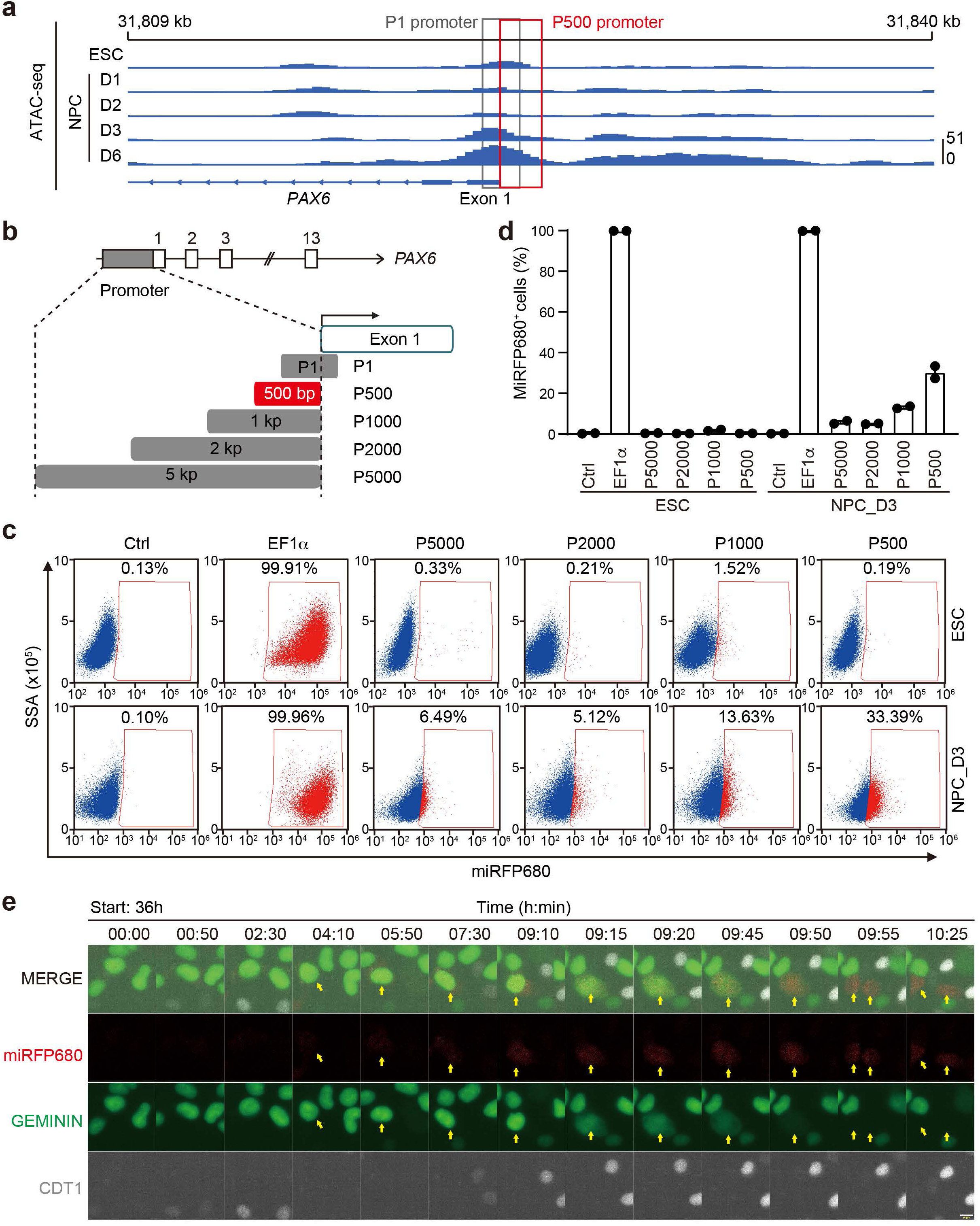
| A novel *PAX6* promoter activates initial *PAX6* expression during the ESC–NPC transition. **a**, ATAC-seq tracks showing chromatin accessibilities at the *PAX6* locus of ESCs and NPCs from D1 to D6 of neural induction. The P1 promoter and the newly identified 500-bp promoter (P500) of *PAX6* are highlighted with grey and red rectangles, respectively. **b**, Schematic diagram of the P1 and candidate *PAX6* promoters tested in reporter assays. **c**, FACS plots quantifying the percentage of cells with miRFP680 expression driven by indicated *PAX6* promoters in ESCs and NPCs at D3 of neural induction (NPC_D3). EF1α was used as the positive control and uninfected cells were used as the negative control (Ctrl). **d**, Quantification of the percentage of miRFP680^+^ cells in **c**. Mean ± range are shown; n=2 independent experiments. **e**, Live-cell imaging of FUCCI NPCs expressing the miRFP680 reporter (red) driven by the P500 promoter during 36–46 hours of neural induction. Cell cycle status is indicated by the FUCCI reporter system (GEMININ, green; CDT1, white). Yellow arrows mark miRFP680^+^ cells. Scale bar, 10 μm.

We next engineered a lentiviral reporter system in ESCs, with the expression of the miRFP680 fluorescence protein driven by the P1 promoter or P1-containing upstream regions (1 kb, 2 kb, or 5 kb; termed P1-1kb, P1-2kb, P1-5kb). Surprisingly, all P1-containing constructs showed expression in both ESCs and NPCs (Extended Data Fig. 5c,d). Thus, P1-based promoters could not recapitulate NPC-specific expression of the endogenous *PAX6*. We found that a 192-bp region of the 410-bp P1 promoter overlapped with the first exon of *PAX6*. This region is known to be bound by OCT4 and NANOG in ESCs^30,31^, providing a potential explanation of the untimely reporter expression in ESCs.

We then tested upstream regions (500 bp, 1 kb, 2 kb, 5 kb; termed P500, P1000, P2000, P5000) adjacent to the *PAX6* transcriptional start site (TSS) (Fig. 3b). These reporters showed negligible activity in ESCs but varying levels of activation in NPC_D3, with P500 exhibiting the highest activity (Fig. 3c,d). To determine if the P500-driven reporter expression was cell cycle-dependent, we introduced the P500 reporter into FUCCI-expressing ESCs and simultaneously tracked its expression and cell cycle status during neural induction. Live-cell imaging analysis identified 90 cells exhibiting miRFP680 reporter activation, all of which initiated reporter expression at G2 (Fig. 3e), mirroring endogenous PAX6. The P500-driven miRFP680 reporter was not expressed in 293FT cells (Extended Data Fig. 5e), suggesting regulation by neural induction-specific transcription factors. Together, these results identify the P500 promoter as the functional promoter driving cell cycle-coupled *PAX6* transcription during NPC specification.

### G2-coupled transcription of *PAX6* is independent of the MMB-FOXM1 complex

The MuvB core component of the DREAM complex interacts with BMYB and FOXM1 to form the MMB-FOXM1 complex, which coordinates gene expression during S and G2/M phases^14^. We investigated whether the MMB-FOXM1 complex contributed to G2-phase-specific PAX6 transcriptional activation. Most G2-expressed genes, such as *CCNB1*, contain cell cycle-dependent elements (CDEs) or cell cycle gene homology regions (CHRs) in their promoters^16^. Analysis of the *PAX6* P500 promoter revealed a putative CHR site predicted to bind LIN54 and FOXM1 (Fig. 4a).

**Fig. 4.**
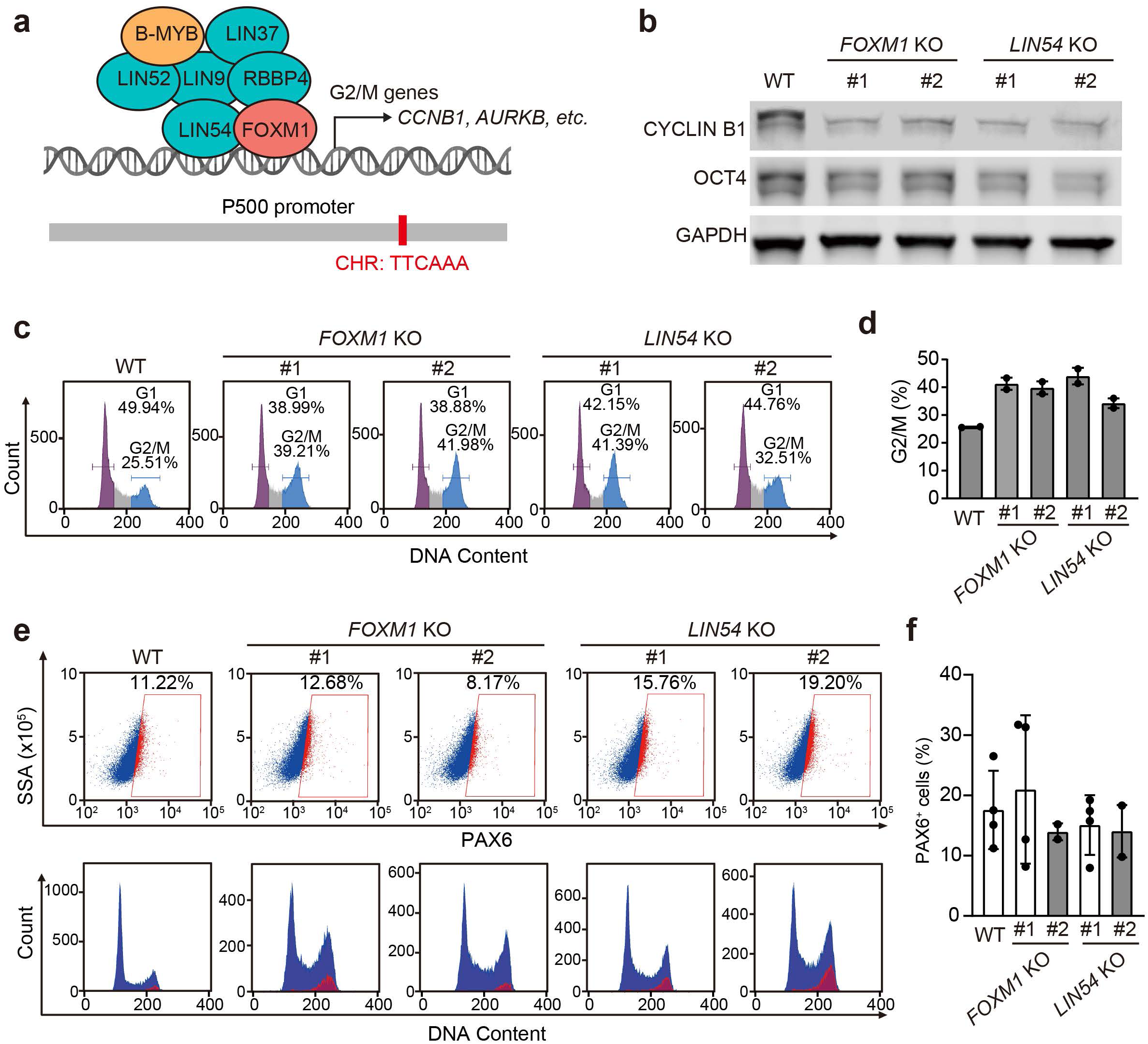
| The DREAM/MMB components FOXM1 and LIN54 are not required for initial *PAX6* expression during ESC–NPC transition. **a**, Schematic illustration of the MMB-FOXM1 complex that drives G2/M-specific gene expression. The *PAX6* P500 promoter contains a putative CHR (cell cycle genes homology region) motif that may be bound by FOXM1 and LIN54. **b**, Western blots showing the protein levels of CYCLIN B1 and OCT4 in wild type (WT), *FOXM1* KO, and *LIN54* KO ESC clones. GAPDH was used as the loading control. **c**, FACS analysis of the cell cycle status of WT, *FOXM1* KO, or *LIN54* KO ESCs. The percentages of cells in G1 and G2/M are indicated. **d**, Quantification of G2/M percentages of cells in **c**. Mean ± range; n = 2 independent experiments. **e**, FACS analysis of PAX6 expression (upper panels) and the cell cycle status (lower panels) of WT, *FOXM1* KO, and *LIN54* KO NPCs at 56 hours of neural induction. **f**, Quantification of PAX6^+^ cells in **e**. n=4 independent experiments for WT, *FOXM1* KO clone #1, and *LIN54* KO clone #1. n=2 independent experiments for *FOXM1* KO clone #2 and *LIN54* KO clone #2.

To assess the potential role of these factors, we used CRISPR-Cas9 technology to knock out *FOXM1* and *LIN54* in ESCs. Two *FOXM1* homozygous knockout (KO) clones were successfully generated using two different sgRNAs (Extended Data Fig. 6a). In contrast, multiple attempts to knockout *LIN54* with three different sgRNAs yielded clones with frameshift mutations in one allele and non-frameshift mutations in the other, suggesting that *LIN54* might be essential for ESC viability and could not be fully deleted (Extended Data Fig. 6b). Despite this, all KO clones exhibited reduced cyclin B1 levels (Fig. 4b), and FACS analysis revealed aberrant cell cycle profiles in *FOXM1*-or *LIN54*-deficient cells, with increased G2/M populations (Fig. 4c,d), confirming their roles in G2-phase gene regulation and mitotic entry.

We next assessed whether *FOXM1* or *LIN54* inactivation influenced PAX6 expression during NPC fate transition. In both *FOXM1* KO and *LIN54* KO lines, the percentage of PAX6^+^ cells in NPC_D2 was not substantially altered, with PAX6 initial expression still occurring in G2 (Fig. 4e,f). Thus, *PAX6* transcriptional activation in G2 appears to be independent of key components of the MMB-FOXM1 complex.

### A 12-bp GC-rich motif is required for *PAX6* transcription during NPC cell fate transition

We next investigated the molecular mechanism driving *PAX6* expression during G2. To identify critical regulatory elements, we generated a series of 50-bp truncations within the P500 promoter (designated P500_Δ1 to Δ10) and delivered these constructs into ESCs using lentiviruses (Fig. 5a). FACS analysis revealed that none of the reporter constructs were expressed in undifferentiated ESCs. Upon differentiation into NPCs, the intact P500 promoter robustly activated the expression of miRFP680. Fluorescence signals were, however, significantly reduced in cells carrying the P500_Δ2 promoter (Fig. 5b,c), implicating the second 50-bp region in transcriptional activation. Consistent with our findings that the MMB-FOXM1 complex components were not required for *PAX6* expression, deletion of the 8th 50-bp region, which contained the putative CHR motif, did not reduce the expression of PAX6 during NPC formation.

**Fig. 5.**
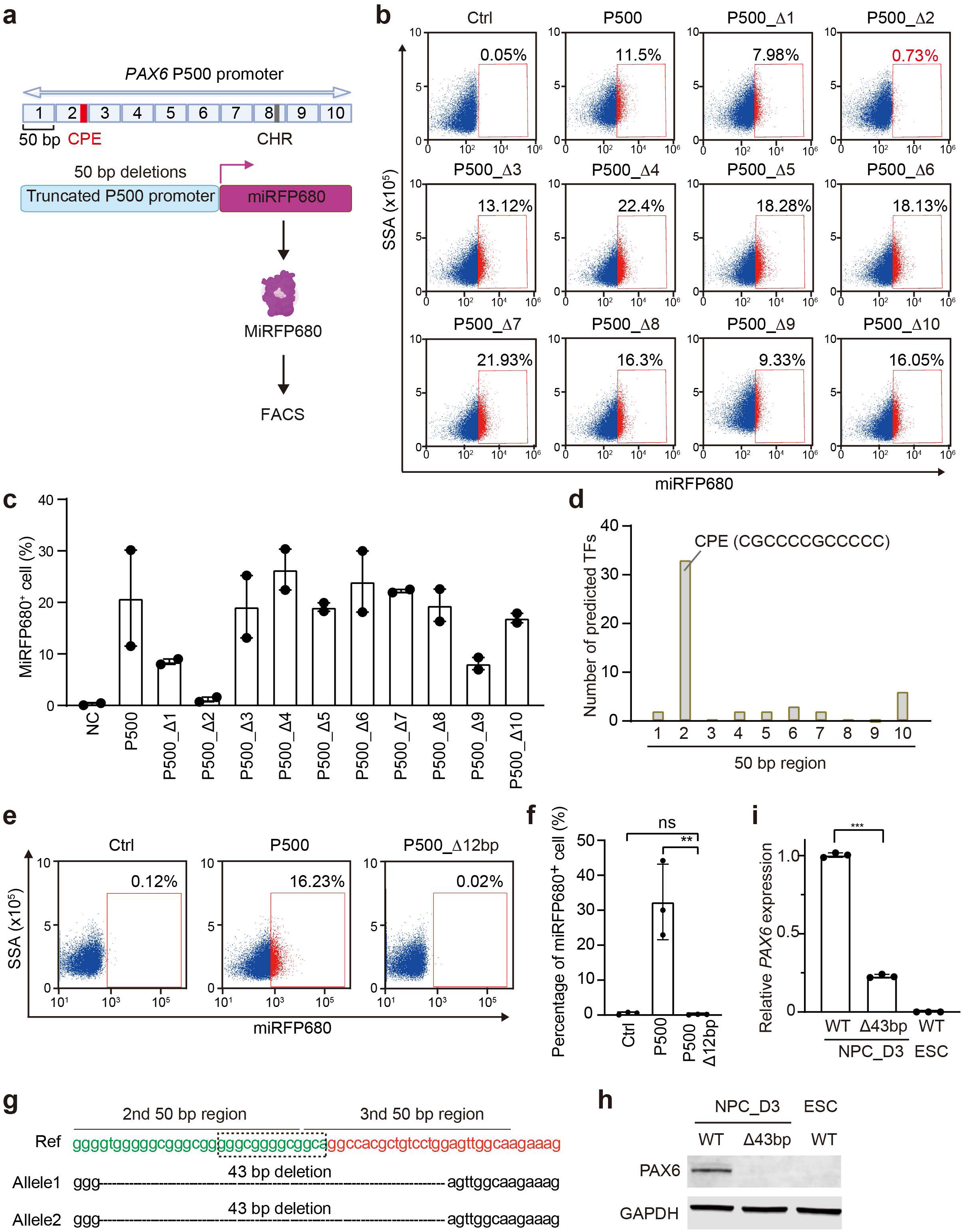
| The core promoter region (CPE) of the P500 promoter is required for initial *PAX6* expression during the ESC–NPC transition. **a**, Schematic illustration of the miRFP680 reporter assay driven by truncated P500 promoters in NPCs. The P500 promoter was divided into ten 50-bp segments. Each fragment was deleted from the P500 promoter to generate a series of truncated promoters termed P500_Δ1, P500_Δ2, P500_Δ3, and so on. The core promoter element (CPE) identified in this study and the putative CHR motif are indicated. **b**, FACS analysis of the percentage of miRFP680^+^ cells in NPCs at day 3 of neural induction. The miRFP680 reporter was driven by the truncated P500 promoters described in **a**. Uninfected cells were used as the negative control (Ctrl). **c**, Quantification of miRFP680^+^ cells in **b**. Mean ± range; n=2 independent experiments. **d**, Distribution of the top 50 predicted transcription factors (TFs) in each 50-bp segment of the P500 promoter. The TF-binding consensus sequence in the 2nd segment is shown. **e**, FACS analysis of miRFP680 expression in NPCs expressing the miRFP680 reporter driven by the P500 promoter with the 12-bp CPE deleted (P500_Δ12bp) at day 3 of neural induction. **f**, Quantification of the percentage of miRFP680^+^ cells in **e**. n=3 independent experiments; ns: p>0.05, **p (0.001, Student’s unpaired two-tailed t-test. **g**, Sanger sequencing results showing the 43-bp deletion in the endogenous *PAX6* promoter in the Δ43bp ESC line. DNA sequences of the 2nd and 3rd 50-bp segments of the P500 promoter are in green and red fonts, respectively. The 12-bp CPE is boxed. **h**, Western blots showing the expression of PAX6 protein in ESCs and NPCs at day 3 of neural induction (NPC_D3). NPCs were derived from WT ESCs or ESCs with the 43-bp deletion in the endogenous *PAX6* promoter. **i**, Quantification of PAX6 protein levels in **h**. Mean ± SD; n=3 independent experiments.

Using the JASPAR database, we predicted transcription factor (TF) binding sites within the P500 promoter (Extended Data Table 1). Interestingly, 33 of the top 50 predicted TFs targeted a 12-bp GC-rich sequence within the second 50-bp region (Fig. 5d), suggesting this motif as a core promoter element regulating *PAX6* transcription. Indeed, the P500_Δ12bp promoter lacking the 12-bp sequence could not activate the transcription of the reporter in NPC_D3 (Fig. 5e,f).

We next attempted to delete the 12-bp motif from the endogenous *PAX6* promoter using CRISPR/Cas9. Due to technical challenges posed by the high GC content surrounding this motif, we could only obtain an ESC line with a 43-bp deletion spanning this region on both alleles (Δ43bp ESCs; Fig. 5g). While Δ43bp ESCs maintained pluripotency marker expression and normal morphology (Extended Data Fig. 6c), PAX6 induction was severely impaired upon differentiation to NPC_D3 (Fig. 5h,i). These results identify a novel GC-rich core promoter element (CPE) essential for *PAX6* transcription during NPC fate transition.

We next sought to identify key TFs regulating PAX6 transcriptional initiation during G2 in NPC fate transition. Using the JASPAR database, we selected the top five predicted TFs that bind the CPE for analysis (Extended Data Fig. 7a). *EGR1* exhibited a temporal expression pattern during ESC–NPC differentiation that closely mirrored *PAX6* (Extended Data Fig. 7b). Single-cell RNA-seq further showed *EGR1* transcript enrichment in G2/M (Extended Data Fig. 7c). The EGR1 protein was present in ESCs and throughout NPC differentiation (up to NPC_D6; Extended Data Fig. 7d). We thus tested whether EGR1 regulated *PAX6* transcription. Using CRISPR-Cas9, we generated *EGR1* KO ESC lines (Extended Data Fig. 7e). After three days of neural induction, *EGR1* KO clones showed undetectable EGR1 protein, but the protein levels of PAX6 were not substantially altered (Extended Data Fig. 7f,g), indicating that EGR1 was not required for PAX6 expression. Individual deletion of other predicted TFs—including PATZ1, ZNF454, and SP8—also had no effect on cell cycle-dependent *PAX6* expression during NPC fate transition. Thus, PAX6 expression might be regulated by a novel transcription factor or by the redundant actions of the tested transcription factors.

### The G2 phase is essential for NPC cell fate transition

To investigate the biological significance of G2-specific transcriptional activation in NPC fate transition, we blocked the cell cycle prior to the G2 phase during differentiation using hydroxyurea (HU), a DNA synthesis inhibitor that arrested cells in S phase. We administered 250 μM HU to the culture medium starting on D1 or D2 of differentiation, with treatment durations of 48 or 24 hours, respectively, culminating at D3 (Fig. 6a). FACS analysis revealed that DMSO-treated control cells maintained normal cell cycle progression, with 55.2% cells expressing PAX6 by D3 (Fig. 6b,c). In contrast, HU-treated cells exhibited prolonged S-phase arrest and a complete absence of PAX6 expression (Fig. 6b,c). IF analysis further demonstrated that HU-treated cells failed to express PAX6 by D3 and displayed enlarged nuclei compared to controls (Fig. 6d). These findings indicate that S-phase arrest disrupts normal NPC fate transition, either by preventing progression along the same trajectory or by altering differentiation trajectories.

**Fig. 6.**
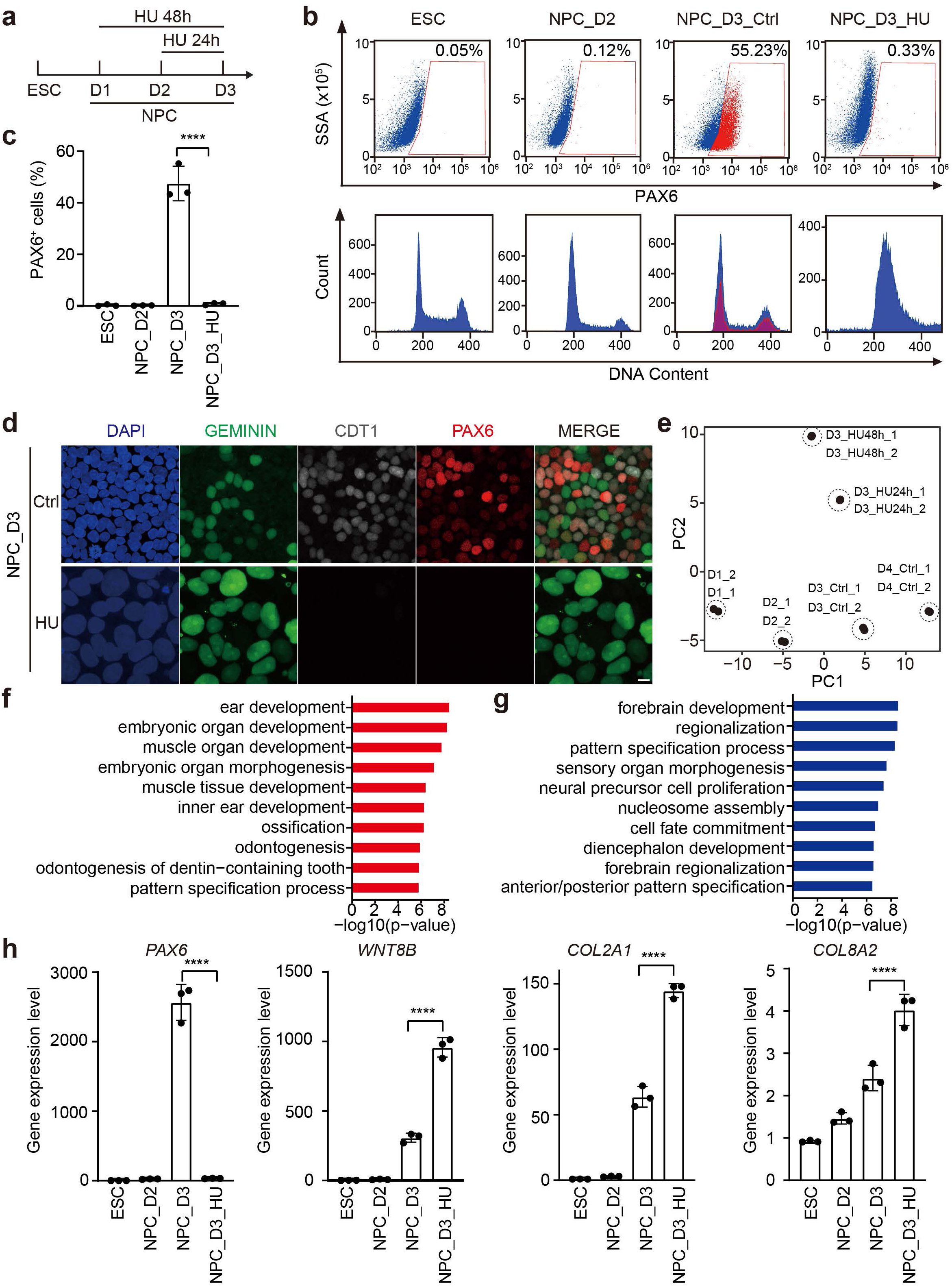
| S phase arrest blocks *PAX6* expression and ESC–NPC transition. **a**, Experimental scheme of hydroxyurea (HU) treatment during neural induction. **b**, FACS analysis of PAX6 expression (upper panels) and cell cycle status (lower panels) of ESCs and NPCs at day 2 (NPC_D2) or day 3 (NPC_D3) of neural induction, with DMSO (Ctrl) or HU treatment. **c**, Quantification of the percentage of PAX6^+^ cells in **b**. Mean ± SD; n=3 independent experiments. **d**, Images of FUCCI NPCs at D3 of neural induction with or without HU treatment stained with DAPI and the PAX6 antibody (red). The cell cycle status was determined by FUCCI reporters (GEMININ, green; CDT1, white). Scale bar, 10 μm. **e**, Principal component analysis (PCA) of transcriptomic profiles of cells at various differentiation stages, with or without HU treatment. **f**,**g**, Gene Ontology (GO) enrichment analysis of upregulated (**f**) and downregulated (**g**) DEGs in HU-treated NPC_D3 cells compared to DMSO-treated NPC_D3 cells. Top ten pathways are shown. **h**, Quantitative PCR analysis of the relative expression of PAX6 and representative mesoderm-related genes in ESCs, NPC_D2, and NPC_D3 with or without HU treatment. Mean ± SD; n=3 independent experiments. **** P<0.0001, Student’s unpaired two-tailed t-test.

To distinguish between these two possibilities, we performed transcriptomic profiling across differentiation stages with or without HU treatment. Principal component analysis (PCA) revealed a clear trajectory for untreated cells from D1 to D4, whereas HU-treated cells—regardless of treatment duration—clustered distinctly from D3 controls. 48-hour HU treatment induced greater divergence than 24-hour treatment, suggesting that cell cycle arrest fundamentally perturbs differentiation rather than merely delaying it (Fig. 6e).

Transcriptome comparison of D3 HU-treated (24-hour duration) and untreated cells (NPC_D2 or NPC_D3) highlighted divergent pathways. Control differentiating NPCs upregulated genes enriched in neurodevelopmental pathways (Extended Data Fig. 8a), whereas HU-treated cells activated mesenchymal lineage programs, including urogenital, renal, and muscular systems (Extended Data Fig. 8b). Direct comparison of treated versus untreated D3 cells confirmed upregulation of mesenchymal signatures (Fig. 6f) and suppression of neurodevelopmental pathways (Fig. 6g). qPCR analysis confirmed significant downregulation of *PAX6* and upregulation of mesoderm-associated genes (*WNT8B*, *COL2A1*, *COL8A2*) in HU-treated cells (Fig. 6h).

Taken together, these results suggest a requirement for G2 phase progression in NPC fate transition. S-phase arrest not only halts neural differentiation but also redirects cells toward mesenchymal lineages, underscoring the critical role of cell cycle dynamics in dictating progenitor cell fate.

## Discussion

In this study, using an ESC–NPC *in vitro* differentiation system and PAX6 as the NPC fate marker, we demonstrate that the transition from ESCs to NPCs occurs specifically during the G2 phase. Our findings further suggest that the G2 phase is required for successful NPC fate determination, highlighting the critical role of the cell cycle in lineage commitment. Mechanistically, we identify a novel 500 bp *PAX6* promoter (P500) responsible for cell cycle-dependent transcriptional activation. Interestingly, the previously reported *PAX6* P1 promoter in neural tissues drives untimely *PAX6* expression in ESCs, suggesting the existence of negative regulators in ESCs that act through the P500 promoter. Systematic dissection of the P500 promoter revealed a core promoter element (CPE) essential for initiating *PAX6* expression. Thus, *PAX6* expression during NPC fate specification is controlled by both negative and positive regulators, whose availability or functions may vary across cell cycle stages (Fig. 7). Bioinformatic predictions and candidate approaches (e.g. DREAM and MMB-FOXM1 complexes) have so far failed to identify TFs driving this process. We note that, while it is relatively straightforward to identify functional cis-regulatory elements in gene regulation, it is notoriously difficult to identity TFs that act through these elements. In addition to *PAX6*, several other genes initiate their expression in G2 during neural induction. It will be interesting to test whether their expression is activated by a similar mechanism.

**Fig. 7.**
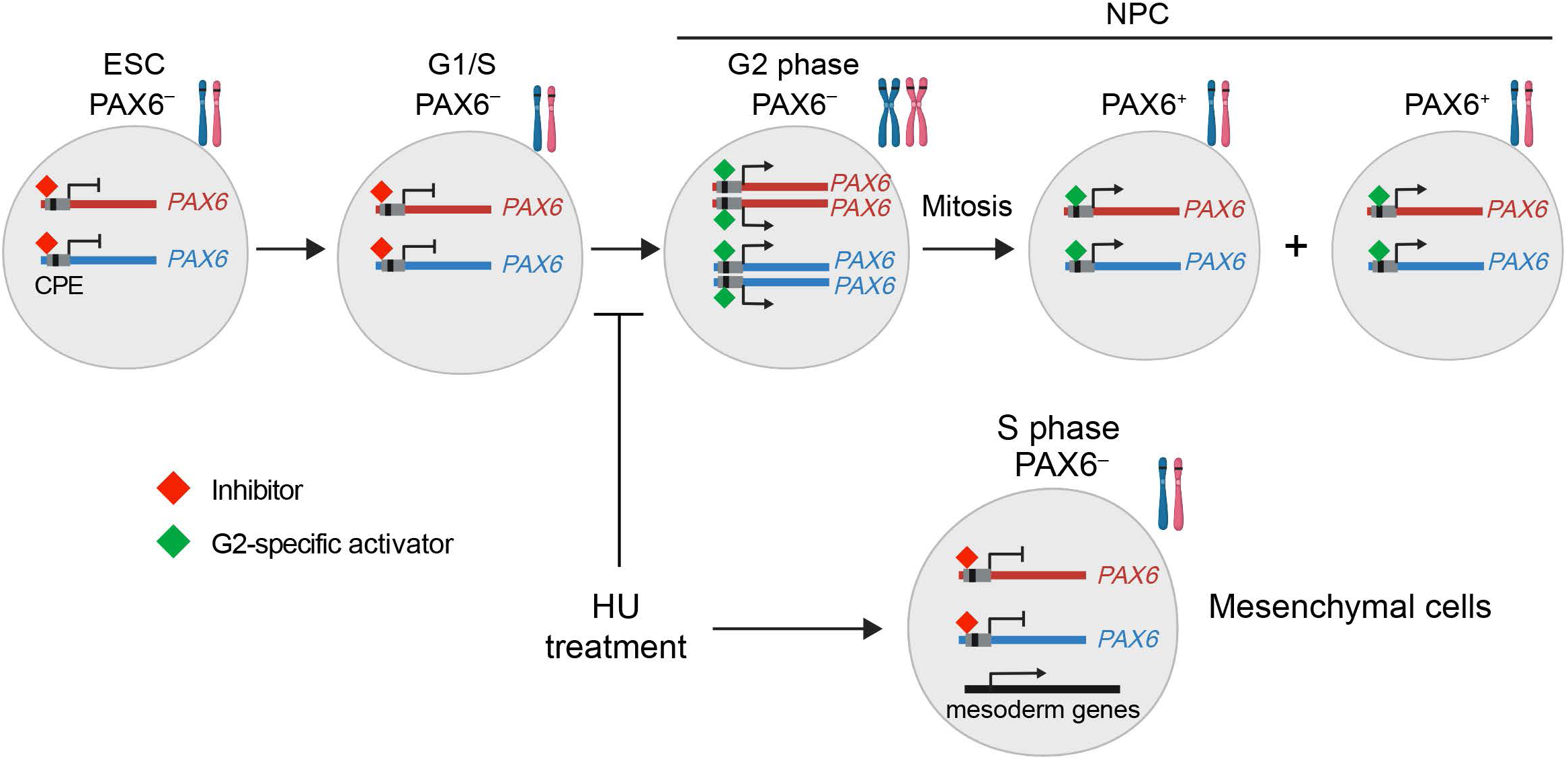
| Model of G2-coupled *PAX6* activation during ESC–NPC transition. *PAX6* transcription is controlled by both negative and positive regulators that act through the P500 promoter. The inhibitors exist in ESCs and at G1 and S phases of cells undergoing the ESC– NPC transition. G2-specific activators that bind the core promoter element (CPE) promote *PAX6* expression at G2. This G2-coupled activation of *PAX6* expression allows concerted activation of all four copies of the *PAX6* gene and generation of two daughter cells of identical fate. Blocking the cell cycle at S phase by hydroxyurea (HU) during differentiation prevents PAX6 activation and leads to aberrant expression of mesoderm genes.

During early human embryogenesis, PAX6 orchestrates neural plate patterning and maintains the undifferentiated state of neuroepithelial cells^32^. Symmetric divisions of PAX6^+^ progenitors ensure rapid expansion of the progenitor pool prior to cortical specialization. Our study reveals that *PAX6* transcription initiates in G2, with subsequent mitosis producing two PAX6^+^ daughter cells. This G2-specific activation may facilitate symmetric divisions, enabling efficient progenitor pool expansion—a process critical for proper CNS development.

Prior studies have emphasized the G1 phase as a pivotal window for fate regulation, showing correlations between G1 lengthening and differentiation onset^33^ ^20,21,34^. By combining live-cell imaging, FUCCI reporters, and scRNA-seq, we instead pinpointed *PAX6* activation to the G2 phase. We speculate that G2-phase fate transitions confer certain advantages: chromatin reorganization and transcriptional reprogramming in G1/S phases risk errors during DNA replication, whereas G2-phase commitment immediately precedes mitosis, ensuring faithful inheritance of fate-determining factors by daughter cells (Figure 7). This mechanism may be particularly beneficial during early development when the expansion of progenitor pools is critical. It will be interestingly to examine whether G2-specific fate commitment occurs in other cell lineages that undergo rapid expansion during development.

Finally, we have shown that S-phase arrest by hydroxyurea (HU) disrupts NPC fate transition and activates mesodermal genes, underscoring the role of G2 phase in neural lineage specification (Figure 7). Our results seemingly contradict a recent study by Kukreja *et al.*, which reported that zebrafish embryos treated with HU and aphidicolin completed germ layer differentiation despite the cell cycle blockade^35^. This apparent contradiction may reflect differences between species or stem from system-specific compensatory mechanisms. In zebrafish, signaling networks (e.g. Notch and Wnt) may buffer fate determination^36,37^, whereas *in vitro* models lack such feedback loops, heightening sensitivity to cell cycle perturbations. Furthermore, HU treatment timing—aligned with early stage of gastrulation in zebrafish versus early neural induction in our system—may differentially impact fate commitment. Future studies should explore how cell-cell communications and microenvironmental cues modulate cell cycle-fate coupling across species, offering insights into developmental robustness.

## Conclusion

Using PAX6 as a fate transition marker in an *in vitro* differentiation system, we provide the first direct evidence that early neural commitment of human ESCs occurs during the G2 phase. We identify a 500 bp *PAX6* promoter governing its transcriptional dynamics and define a core promoter element (CPE) essential for activation. We further show that S-phase arrest blocks NPC specification by diverting cells toward mesodermal lineages, implicating an instructive role of the cell cycle in fate determination. These findings advance our understanding of how cell cycle dynamics integrate with transcriptional programs to orchestrate lineage transitions during development.

## Methods

### Human embryonic stem cell (ESC) culture

Human embryonic stem cell line (ESC; H9/WA09) was obtained from WiCell Research Institute, Wisconsin, USA. ESCs were cultured in mTeSR1 media (STEM CELL Technology, #85850) in 6-well plates coated with Matrigel (Corning, #354277)^38^. ESCs were passaged to new plates when they grew to ∼70% confluency. The Versene reagent (Thermo Fisher Scientific, #15040066) was used for routine passage, and 10 μM Y-27632 (Stem Cell Technologies, #72308) was added for 24 hours to prevent cell apoptosis. All ESCs were cultured at 37°C in a humidified atmosphere supplemented with 5% CO_2_. The pluripotency of ESCs was confirmed by immunofluorescence staining of pluripotency markers, such as OCT4, SOX2, and NANOG. ESCs were regularly tested for mycoplasma contamination by using a commercial kit (MycoBlue Mycoplasma Detector, Vazyme, D101-02).

### Neural progenitor cell (NPC) differentiation and hydroxyurea treatment

ESCs were differentiated into NPCs using a monolayer protocol with the STEMdiff^TM^ SMADi Neural Induction Kit (Stem Cell Technologies, #08581) according to the manufacturer’s instructions^22^. Briefly, ESCs cultured in mTeSR1 were detached into single-cell suspension using Accutase (Stem Cell Technologies, #07920), and cells were resuspended in the neural induction medium containing SMAD inhibitor supplements. 2 × 10^6^ cells were seeded into Matrigel-coated 6-well plate in the neural induction medium with SMAD inhibitors, supplemented with 10 μM Y-27632. The cells were replenished with fresh neural induction medium containing SMAD inhibitors every day and passaged to new plates every 6 days. 10 μM Y-27632 was added for 24 hours to prevent cell apoptosis after each cell passaging.

For cell cycle synchronization at the S phase, 250 μM hydroxyurea (HU) was added to the neural induction medium from day 1 (D1) or D2 of differentiation, and these cells were cultured for another 48 or 24 h, respectively, until D3 of differentiation before analysis of PAX6 expression.

### Forebrain neuron differentiation

For forebrain-type neuron (FBN) differentiation, the ESC-derived mature NPCs (passage 3; D18) were detached using Accutase and seeded onto a Matrigel-coated 6-well plate at a density of 1.25 × 10^5^ cells/cm^2^ in STEMdiff Neural Induction Medium containing SMAD inhibitors for 24 h. The medium was then changed to STEMdiff Forebrain Neuron Differentiation medium (Stem Cell Technologies, #08600) and replenished with fresh medium daily for 6 days. On D7, forebrain neural precursors were detached using Accutase and seeded onto a Matrigel-coated 6-well plate at a density of 5 × 10^4^ cells/cm^2^ in STEMdiff Forebrain Neuron Maturation medium (Stem Cell Technologies, #08605)^39^, with a full-medium change every 2 d for 15 d.

### Plasmids

For gene editing, the sgRNAs targeting the gene of interest were subcloned into the pLenti-CRISPR v2 vector (Addgene, #5296), and the sgRNA targeting the *AAVS1* site was used as a negative control. All sgRNA sequences were listed in Supplementary Table 1. For the *PAX6* reporter assay, the promoters of *PAX6* with different lengths were amplified from the genomic DNA of ESCs and ligated with cDNAs encoding the miRFP680 reporter in a lentiviral plasmid backbone. The ClonExpress MultiS One Step Cloning Kit (Vazyme, #C113-01) and Mut Express II Fast Mutagenesis Kit V2 (Vazyme, #C214-01) were used to generate these constructs.

### Lentivirus packaging and generation of FUCCI ESCs

The lentiviral targeting vectors were transfected into HEK293FT cells along with psPAX2 (Addgene, #12260) and pMD2.G (Addgene, #12259) using Lipofectamine 2000 (Life Technologies, #11668019). Following transfection, viral supernatant was collected at 2-3 days and concentrated using a Lenti-X concentrator (Takara, #PT4421-2).

The FUCCI vector was obtained from Addgene (#86849) and used for generating lentiviruses. ESCs were infected with the FUCCI lentivirus and then treated with 0.5 μg/ml puromycin for 3 d. The surviving cells were picked as single clones to establish the FUCCI-ESC cell line. FUCCI signals and pluripotency markers of the FUCCI ESCs were verified by immunofluorescence staining.

### Western blotting

The cell pellets were lysed in 1×RIPA buffer containing 1 mM PMSF protease inhibitor and genomic DNA was sheared by sonication. The resulting lysates were cleared through centrifugation at 4°C, analyzed by SDS-PAGE, and then transferred to the PVDF membrane using the wet-transfer protocol. The primary antibodies used in this study included: rabbit anti-PAX6 (Cell Signaling Technology, #60433); rabbit anti-GAPDH (Proteintech, #60004-1-Ig); mouse anti-OCT3/4 (Cell Signaling Technology, #2750S). The secondary antibodies used were donkey anti-rabbit IgG (H + L) Dylight 680 conjugates (Cell Signaling Technology, #5366S) and donkey anti-mouse IgG (H + L) Dylight 800 conjugates (Cell Signaling Technology, #5257S). The membranes were scanned and quantified using the Odyssey infrared imaging system (LI-COR Biosciences).

### Quantitative real-time PCR

Total RNAs were extracted from ESCs, NPCs, or FBNs using the FastPure Cell/Tissue Total RNA Isolation Kit V2 (Vazyme, #RC112-01), reversely transcribed and amplified to prepare the long ssDNA using the Guide-it long ssDNA kit (Takara, #632667) following the manufacturer’s instructions. Real-time qPCR was performed by using iTaq™ Universal SYBR® Green Supermix (Bio-Rad, #1725124) in a Bio-Rad CFX96 system, and the primers used in this study were listed in Supplementary Table 1.

### Immunofluorescence staining

ESCs, NPCs, and FBNs were cultured on Matrigel-coated cover glasses (Thermo Fisher Scientific, #1254580). The cells were briefly rinsed with 1×PBS, then fixed in 4% paraformaldehyde solution for 15 min at room temperature. The cells were permeabilized in PBS containing 0.2% Triton for 10 min and then incubated in the blocking buffer consisting of 3% BSA and 0.1% Tween 20 in PBS for 1 h at room temperature. After blocking, the cells were incubated with primary antibodies diluted in the blocking buffer for 2 h at room temperature. The cells were subsequently washed three times with the wash buffer consisting of 0.1% Tween 20 in PBS, then incubated with secondary antibodies diluted in the blocking buffer for 1 h at room temperature. After another three washes in the wash buffer, the cells were co-stained with 1 μg/ml DAPI. The cover glasses were then mounted on glass slides using the Fluoromount-G® Mounting Medium (SouthernBiotech, #0100-01) and sealed with nail polish. Slides were imaged using a 40×oil/water objective on a ZEISS 900 confocal microscope. ImageJ was used for image processing and quantification^40^.

The primary or secondary antibodies used for staining included rabbit anti-PAX6 (Cell Signaling Technology, #60433), mouse anti-OCT3/4 (BD Biosciences, #560307), goat anti-NANOG (R&D Systems, AF1997), mouse anti-TUJ1 (R&D Systems, #MAB1195), donkey anti-rabbit IgG (H+L) antibody Alexa Fluor™ 488 (Invitrogen, #A32790), donkey anti-rabbit IgG (H+L) antibody with Alexa Fluor™ 647 (Invitrogen, #A32795), and donkey anti-rabbit IgG (H+L) antibody with Alexa Fluor™ 594 (Invitrogen, #A32754).

### Flow cytometry

The cells were dissociated into single cells with Accutase and briefly washed with PBS. The cell pellets were then fixed with BD Phosflow™ Fix Buffer I (BD Biosciences, #554655) for 20 min at room temperature. After two washes with PBS, the cells were resuspended in BD Phosflow™ Perm Buffer III (BD Biosciences, #558050) and incubated for 30 min on ice or stored at –20°C. For staining with the PAX6 antibody, approximately 5×10^5^ cells stored in BD Phosflow™ Perm Buffer III were washed twice with PBS containing 1% BSA and then incubated with the rabbit anti-PAX6 antibody (Invitrogen, #42-6600) at 1:50 dilution for 2 h at room temperature. After two washes with PBS containing 1% BSA, the cells were incubated with Alexa Fluor-488 conjugated donkey anti-rabbit IgG (H + L) antibody (Invitrogen, #A32790) at a 1:1000 dilution for 1 h in the dark at room temperature. After two additional washes with PBS containing 1% BSA, the cells were resuspended in 100 μl of PBS containing 1 μg/ml DAPI. Finally, the cells were filtered using a 40-μm cell strainer and analyzed on a CytoFLEX LX instrument (Beckman).

### Time-lapse imaging

The differentiating cells were cultured in Matrigel-coated Nunc Lab-Tek chambered cover glass (Thermo Fisher Scientific; cat#155411). Imaging was conducted at 5-min intervals over a period of 30–72 h after neural induction, using a 40×objective on a Delta Vision microscope (GE Healthcare). The microscope was equipped with an environmental chamber regulating temperature and CO_2_ levels. Time-lapse videos were processed and quantified using ImageJ.

### RNA sequencing (RNA-seq)

All bulk RNA-seq experiments were conducted with biological duplicates. Total RNAs from ESCs, NPCs, or FBNs were extracted using the FastPure Cell/Tissue Total RNA Isolation Kit V2 (Vazyme, #RC112-01), and sequencing libraries were generated using the NEBNext Ultra RNA Library Prep Kit for Illumina (NEB, USA, #E7530L) following the manufacturer’s instructions. Briefly, mRNAs were purified from total RNAs using poly-T oligo-attached magnetic beads. Fragmentation was conducted using divalent cations at an elevated temperature in NEB Next First Strand Synthesis Reaction Buffer (5×). First-strand cDNAs were synthesized using random hexamer primers and M-MuLV reverse transcriptase (RNase H). Subsequently, second-strand cDNA synthesis was performed using DNA polymerase I and RNase H. The remaining overhangs were converted into blunt ends through exonuclease/polymerase activities. Following adenylation of the 3’ ends of DNA fragments, NEB Next Adaptors with hairpin loop structures were ligated for hybridization preparation. cDNA fragments within the desired length range of 370∼420 bp were purified using the AMPure XP system (Beverly, USA). 3 µl USER Enzyme (NEB, USA) was incubated with size-selected, adaptor-ligated cDNA at 37°C for 15 min, followed by 5 min at 95°C prior to PCR. PCR was performed using the Phusion high-fidelity DNA polymerase, universal PCR primers, and the index (X) primer. The PCR products were purified using the AMPure XP system, and the library quality was assessed on the Agilent 5400 system (Agilent, USA). The libraries were then quantified using QPCR (1.5 nM) and pooled for sequencing with the PE150 strategy on Illumina platforms from Novogene Bioinformatics Technology Co., Ltd (Beijing, China), based on the effective library concentration and desired data amount.

The quality control of FASTQ files was conducted using FastQC (v0.11.9) and Fastp^41^. The sequences were aligned to the Hg38 human genome build using the STAR alignment tool (v2.7.8a)^42^. The quantification of gene expression was performed using the raw count metrics, based on RefSeq gene annotation, with the aid of FeatureCounts in subread (v2.0.2)^43^. Differential expression analysis was performed using the “DESeq2” (v1.38.3)^44^ package in R. Data visualization was done using the “ggplot2” package in R.

### 4D label-free quantitative proteomics

For protein extraction, the cell samples were ground into powder in liquid nitrogen and transferred to a 5-ml centrifuge tube. Subsequently, 4 volumes of the lysis buffer consisting of 8 M urea and 1X protease inhibitor cocktail were added to the cell powder. The mixture was subjected to sonication for 3 min on ice using a high-intensity ultrasonic processor (Scientz). The resulting solution was centrifugated at 12,000 g at 4°C for 10 min to remove any remaining debris. The supernatant was collected, and the protein concentration was determined using the BCA kit following the manufacturer’s instructions.

For the digestion process, the protein solution was incubated in 5 mM dithiothreitol for 30 min at 56°C. Subsequently, alkylation was performed by introducing 11 mM iodoacetamide and reacting for 15 min at room temperature in the dark. The protein samples were diluted with 100 mM TEAB to decrease the urea concentration to less than 2 M. Trypsin was added to the diluted samples at a trypsin-to-protein mass ratio of 1:50 for the initial digestion overnight, followed by a second 4-h digestion at a trypsin-to-protein mass ratio of 1:100. The resulting peptides were purified and separated from impurities using a C18 solid-phase extraction (SPE) column.

For LC-MS/MS analysis, the tryptic peptides were dissolved in solvent A (a mixture of 0.1% formic acid and 2% acetonitrile in water) and directly loaded onto a homemade reversed-phase analytical column, which had a length of 25 cm and an inner diameter of 100 μm. The mobile phase consisted of solvent A and solvent B (0.1% formic acid in acetonitrile). The peptides were separated using the following gradient: 0-70 min with a linear increase from 9% to 30% of solvent B, 70-82 min with a linear increase from 30% to 40% of solvent B, 82-86 min with a linear increase from 40% to 90% of solvent B, and 86-90 min at a constant concentration of 90% of solvent B. The separation was performed at a constant flow rate of 450 nl/min using an Easy-nLC1000_TOF UHPLC system (Bruker Daltonics).

The MS/MS data obtained were processed using the MaxQuant search engine (version 1.6.15.0). Tandem mass spectra were searched against the Homo_sapiens_9606_SP_20230103.fasta database, which contains 20389 entries, and was concatenated with a reverse decoy and contaminants database. Trypsin/P was specified as the cleavage enzyme, allowing up to 2 missing cleavages. The minimum peptide length was set to 7 amino acids, and the maximum number of modifications per peptide was set to 5. The mass tolerance for precursor ions was set to 20 ppm for both the first search and main search, and the mass tolerance for fragment ions was also set to 20 ppm. Carbamidomethyl on cysteine was specified as a fixed modification, while acetylation on protein N-terminal and oxidation on methionine were specified as variable modifications. To ensure high confidence in the identified results, false discovery rate (FDR) thresholds were adjusted to achieve an FDR of less than 1% for proteins, peptides, and peptide-spectrum matches (PSMs).

Fisher’s exact test was used to determine the significance of the functional enrichment of differentially expressed proteins, with the identified protein as the background. Significant functional terms were identified based on a fold enrichment (log2FC) greater than 0.5 and a p-value less than 0.05.

### Time-series cluster (Mfuzz) analysis

To identify proteins with significant changes during neural induction, the relative expression levels of proteins were first transformed into logarithmic values using a Log2 transformation, and then protein candidates with SD>0.5 were selected. For the Mfuzz analysis, the clustering number (k) was set to 4, and the fuzziness coefficient (m) was set to 2. Visualization of cluster results were performed using the mfuzz.plot2 function in the Mfuzz package^45^.

### ATAC-seq

All ATAC-seq experiments were conducted in biological duplicates using the commercial kit (Active Motif, #53150). Briefly, 100,000 cells were used to isolate nuclei. Following cell lysis in an ice-cold ATAC lysis Buffer, nuclei were incubated in the tagmentation master mix for 30 min at 37°C, and the resulting DNA was purified using DNA purification columns. PCR amplification of the tagmented DNA was then performed to generate libraries with appropriate indexed primers. Following SPRI bead clean-up, the DNA libraries were sequenced on the Illumina NovaSeq platform (PE-150, 50 million reads) at Hangzhou Repugene Technology Co., Ltd.

The quality control of the ATAC-seq FASTQ files was conducted using Cutadapt (v4.0)^46^. The sequencing data were aligned to the hg38 human genome assembly using bwa. Duplicate reads were removed using picard, and only non-duplicate reads in the BAM format were used for subsequent analysis. Peak calling on the nucleosome-free reads was performed with MACS2(v2.2.9)^47^ using the ‘callpeak’ module and the following parameters: –shift –100 –extsize 200. For data normalization and visualization, the BAM files were converted to the bigWig format using “bamCoverage” with CPM in deepTools (v3.5.3)^48^. Heatmaps and average profiles were generated using the “plotHeatmap” and “plotProfile” scripts in deepTools. To annotate the location of the ATAC-Seq peaks in terms of important genomic features, their BED files were assigned to promoters (defined as –2 kb from the transcription start site), introns, intergenic regions, exons, etc. using the “ChIPseeker” package in R(v1.42.1)^49^. Using the “DiffBind” package in R (v3.8.4), peaks with significant differences between groups were selected by applying two criteria: p value < 0.05 and absolute fold change (FC) > 2, and gene region annotation was performed using the “ChIPseeker” package in R.

### Single-cell multiome ATAC+RNA sequencing

The single-cell (sc) RNA-seq and ATAC-seq were performed using a 10×Genomics kit (PN-1000280). Cells were collected at different timepoints, counted using the Countess II FL Automated Cell Counter (Thermo Fisher Scientific), and then lysed on ice for 5 min. Subsequently, 12,000 isolated nuclei were transposed and loaded onto the Chromium Next GEM Chip J (10×Genomics). The chip containing the cells was then loaded onto a Chromium Controller, and the library construction was performed according to the manufacturer’s instructions. Sequencing of the gene expression library was performed on an Illumina Novaseq 6000 platform with the PE150 reading strategy. The scATAC library was sequenced on the Illumina Novaseq 6000 platform with a sequencing depth of at least 25 k read pairs per nucleus with the PE50 reading strategy. Sequencing was performed by Hangzhou Lianchuan Biology Technology Co. Ltd., China.

For the preprocessing of single-cell multiomics data, the raw sequencing data resulting from single-cell multiomics experiments were processed using Cell Ranger ARC 1.0.1 software to perform various analyses related to gene expression and chromatin accessibility. Reference human genome data (GRCh38) for Cell Ranger ARC was downloaded from https://cf.10xgenomics.com/supp/cell-arc/refdata-cellranger-arc-GRCh38-2020-A-2.0.0.tar.gz. Finally, cellranger-arc was used to generate feature counts at the single-cell level for scRNA-seq/ATAC-seq analysis.

For merging multiple single-cell RNA-seq counts matrices from different samples generated by Cell Ranger ARC, the merge function of Seurat (v5.0) in R (4.2.1) was used^50^. Low-quality cells with few genes and redundant mitochondrial genes were filtered out, resulting in the retention of 70,887 high-quality cells for subsequent analysis.

Downstream analysis of RNA-seq was all performed with Seurat (v5.0) in R (v4.2.1). Dimensionality reduction and clustering were performed on all cells. Principal component analysis (PCA) and UMAP algorithms were applied to check clustering of samples from each timepoints. Distributions of NPC markers, including PAX6, among cell clusters were shown by Featureplot in Seurat.

CellCycleScoring function in the Seurat package was used to determine the cell cycle phase of each cell, according to a list of markers of S and G2/M phases^51^. Cells not assigned to S and G2/M phases were annotated as G1 phase. Cell cycle phase and development timepoints were combined to create the more delicate phase-level timepoint labels and assigned to each cell. Comparisons of each marker gene expression among different cell cycle phases, development timepoints or phase-level timepoints were shown by Dotplot in Seurat.

Differential expression gene analysis for cell clusters of phase-level timepoints along the development time axis was detected using FindMarkers in Seurat with the Wilcoxon test^44^. The threshold of differential expression of adjusted p-value is 0.05 and that of log2FoldChange is ±2. Volcano plots of differential expression genes were shown using ggplot2 (v3.5.0). GO annotation (Biological Process) of detected differential expression genes was performed with clusterProfiler package (v4.6.2) and the results were shown with dotplot in the enrichplot package (v1.18.4).

This study only analyzed single-cell transcriptome data. Single-cell ATAC-seq data were not used and will be reported in subsequent studies.

### Gene ontology analysis

In the analysis of the bulk RNA-seq dataset, up– or down-regulated genes were selected based on differential expression analysis in the R package DESeq2 (v1.38.3)^44^ with thresholds of log2FoldChange = 1.5 and adjusted p-value = 0.05. For gene ontology analysis, the up– or down-regulated genes were input into the enrichGO function in the R package clusterProfiler (v4.6.2)^52^ with “Biological Process” as the ontology dataset, “org.Hs.eg.db” as the annotation database, and with threshold cutoff = 0.05. The top 10 gene ontology categories were chosen for visualization based on –log10(adjusted p-value).

Up– or down-regulated genes between cell clusters of the scRNA-seq dataset were selected based on the FindMarkers function from Seurat (v5.1.0) with default parameters. GO enrichment in scRNA-seq was performed by compareCluster in R package clusterProfiler (v4.6.2) with similar parameters as those used in bulk RNA-seq except the threshold cutoff was set at 0.01.

For analyzing ATAC-seq datasets with peak-calling, up– or down-regulated peaks were selected by DESeq2 (v1.38.3) and converted into covered genes by the R package GenomicRanges (v1.50.2) and TxDb.Hsapiens.UCSC.hg38.knownGene (v3.16.0) as the input gene list for GO enrichment. GO enrichment of the ATAC-seq and visualization was performed as described for bulk RNA-seq.

### Quantitation and statistical analysis

GraphPad Prism 7 was used to perform the statistical analysis of Western blot, qPCR, confocal imaging, and flow cytometry data. All data were shown as mean ± SD. The Student’s unpaired two-tailed t-test was applied for significance evaluation between the indicated groups. The one-way analysis of variance (ANOVA) was applied to assess pairwise differences. *p(0.01, **p (0.01, ***p(0.001, ****p(0.0001. The number of biological replicates was indicated in the relevant method sections.

## Data and code availability

The multiomics datasets generated in this study were submitted to NCBI (GSE291903, GSE291905, GSE291907, GSE291908). This study did not generate new codes, and all codes used are available online (https://github.com/Rong-ao/NPC_multiomics.git). Any additional information required to reanalyze the data reported in this paper is available from the lead contact, Hongtao Yu, upon request.

## Acknowledgements

We are grateful to Shang Cai, Yihan Wan, and Qi Hu from School of Life Science, Westlake University, for their advice during the project. We also thank Xilai Ding and other members of the FACS core facility at Westlake University for technical support and the Westlake University High-Performance Computing Center for computational resources. This work was supported by the National Natural Science Foundation of China (Project 32130053 to H.Y.) and the New Cornerstone Science Foundation (to H.Y.).

## Author contributions

Project conception: S.H., S.Q., and H.Y.; Investigation: S.H.; Data analysis: S.H., R.K., Z.S., G.L., and Y.Z; Supervision: S.Q., H.W., Y.Z., L.-L.C., H.Y.; Writing the original draft: S.H.; Reviewing and editing: S.H., S.Q., H.W., L.-L.C., H.Y., with the help of all authors.

## Figure legends

**Supplementary Table 1.**
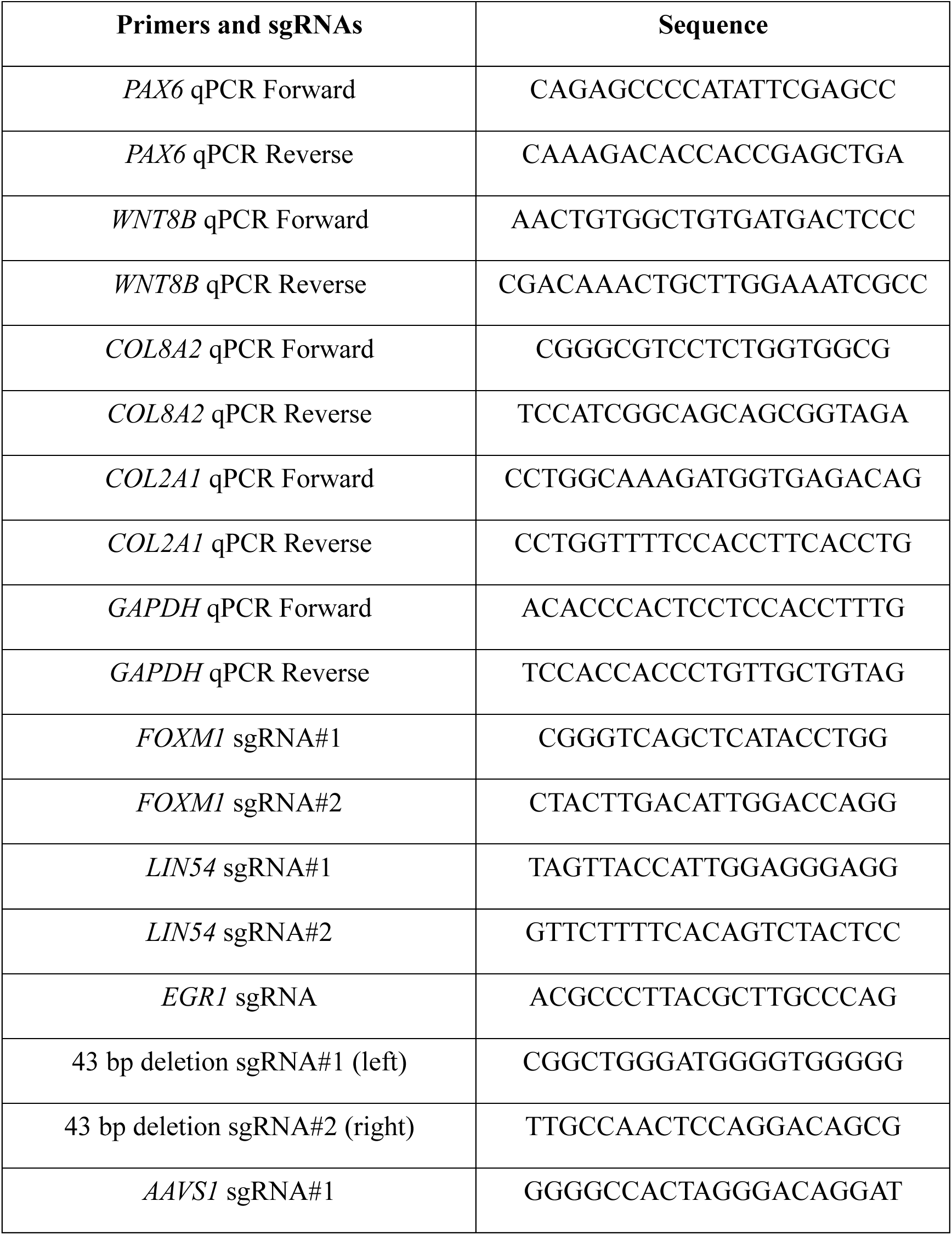
qPCR primers and sgRNA sequences.

**Extended Data Fig. 1.**
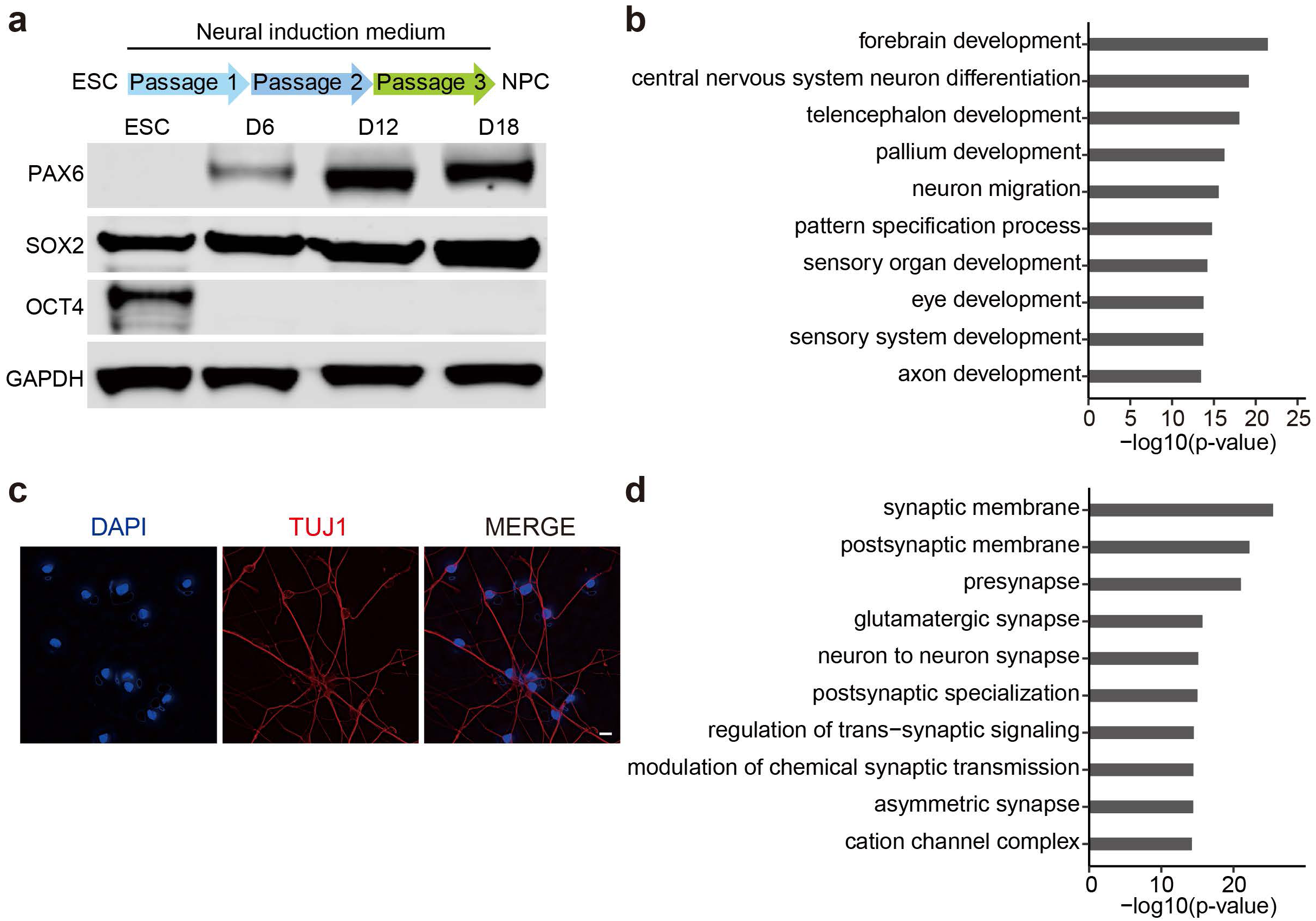
| Establishment and characterization of the in vitro neural induction system from ESCs. **a**, Western blot analysis of the protein levels of PAX6, SOX2, and OCT4 in ESCs and NPCs at day 6 (D6), day 12 (D12), and day 18 (D18) of neural induction. GAPDH was used as the loading control. **b**, Gene Ontology (GO) enrichment analysis of upregulated genes in NPC_D18 compared to ESCs. Top 10 pathways are shown. **c**, Image of forebrain neurons (differentiated from ESC-derived NPCs) stained with DAPI and antibody against the neuron marker TUJ1 (β-III Tubulin). Scale bar, 10 μm. **d**, GO enrichment analysis of upregulated genes of NPC-derived neurons compared to NPC_D18. Top 10 pathways are shown.

**Extended Data Fig. 2.**
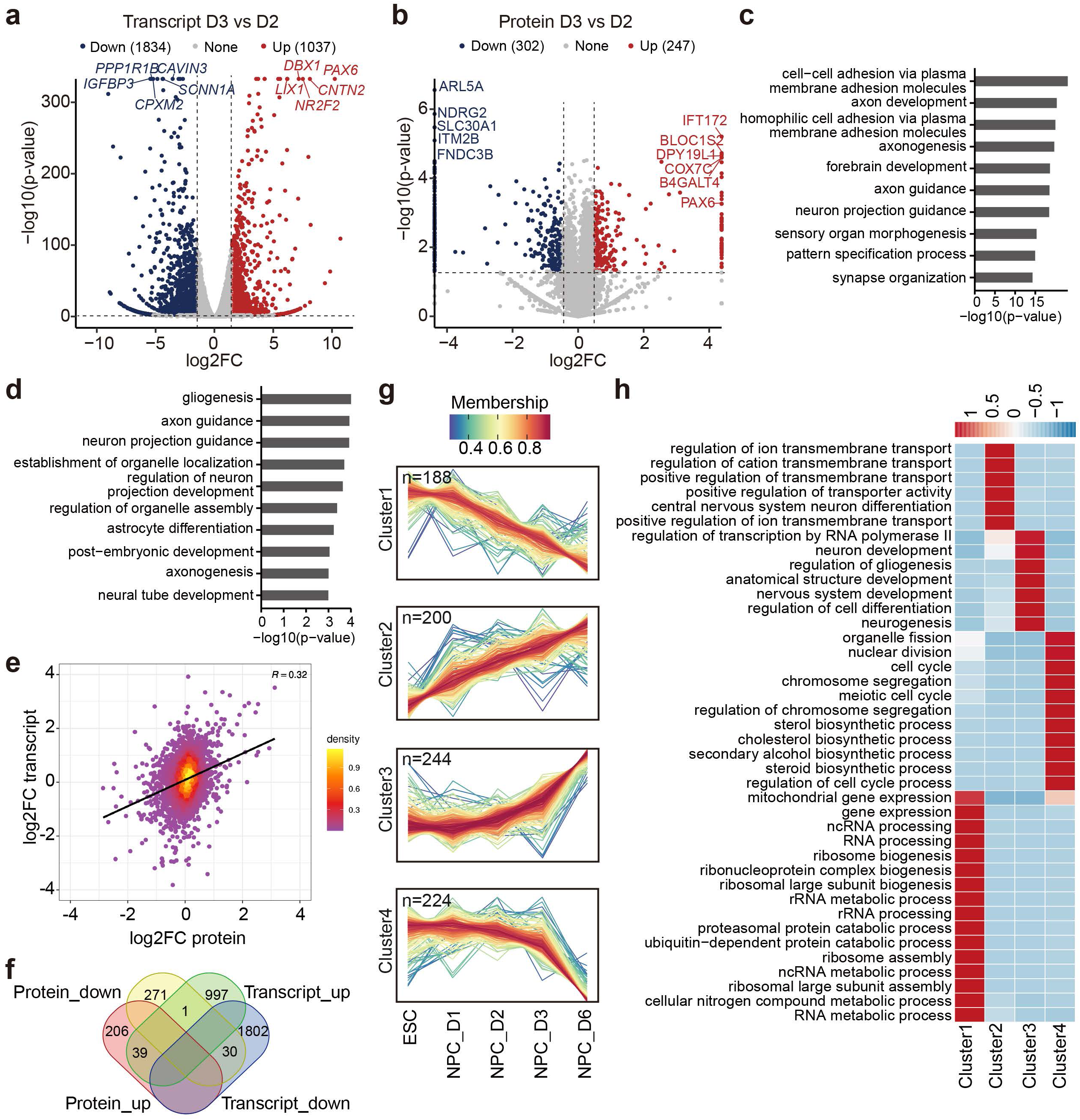
| Transcriptomic and Proteomic characterization of cells during ESC– NPC differentiation. **a**, Volcano plot showing up– and down-regulated genes in NPC_D3 compared to NPC_D2. The horizontal and vertical dotted lines indicate p value(0.01 and |log2FC|>1.5, respectively. **b**, Volcano plot showing up– and down-regulated proteins in NPC_D3 compared to NPC_D2. The horizontal and vertical dotted lines indicate p value(0.05 and |log2FC|>0.5, respectively. **c**, Top 10 enriched GO terms of up-regulated genes in NPC_D3 compared to NPC_D2. **d**, Top 10 enriched GO terms of up-regulated proteins in NPC_D3 compared to NPC_D2. **e**, Scatter plot showing the correlation between the changes in transcripts and the corresponding proteins between NPC_D3 and NPC_D2. The color of the points indicates the density of points at that location. **f**, Venn diagram showing the count numbers of up– and down-regulated transcripts and proteins in NPC_D3 compared to NPC_D2. **g**, Time-series analysis of the proteomics (mfuzz) data of ESCs and NPCs at indicated times of neural induction. **h**, GO analysis of the four protein clusters in **g**.

**Extended Data Fig. 3.**
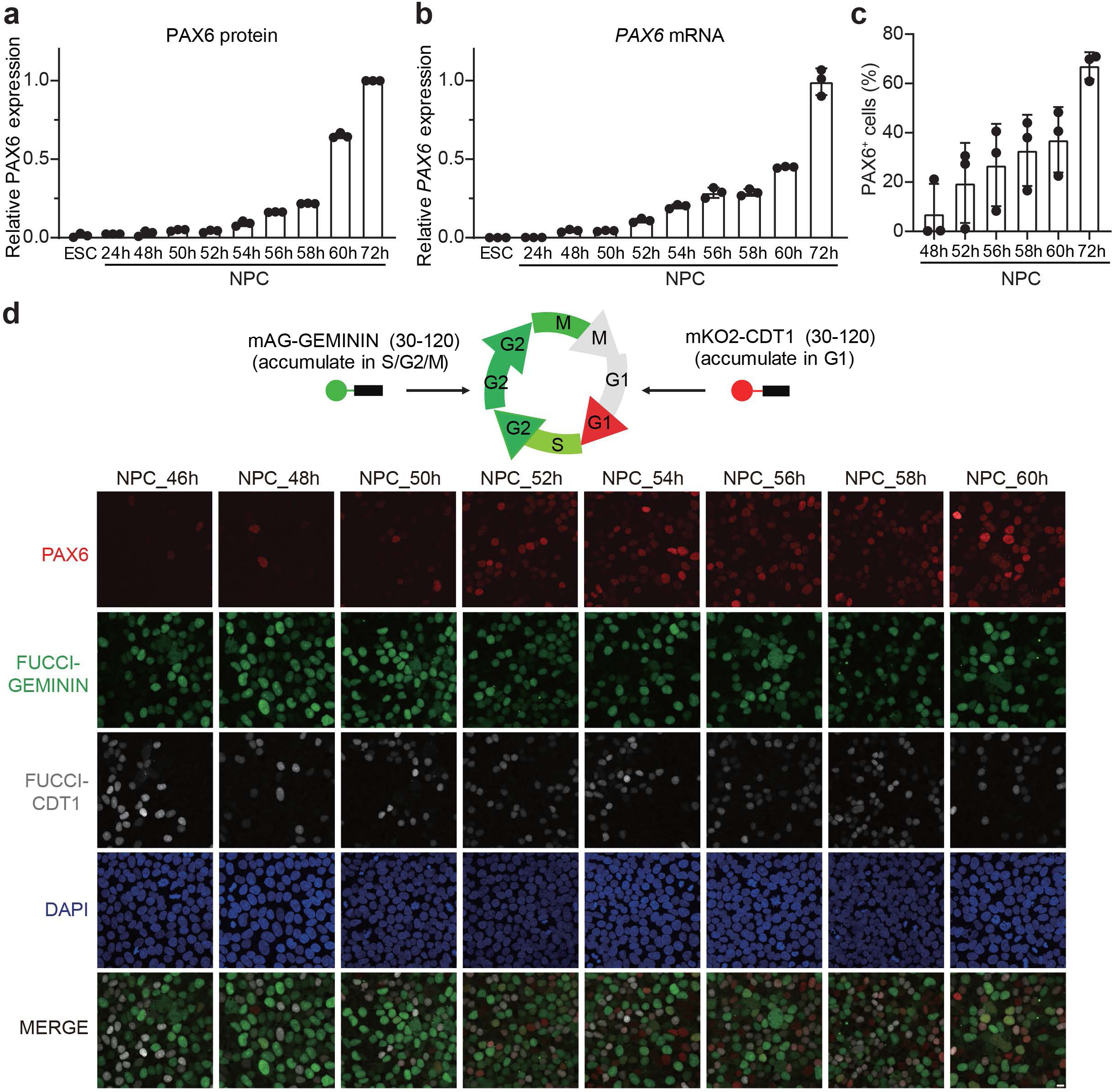
| Initial *PAX6* expression occurs between day 2 and day 3 of neural induction. **a**, Quantification of PAX6 protein levels (detected by Western blot) at different time points during ESC–NPC differentiation. Mean ± SD; n=3 independent experiments. **b**, Quantification of PAX6 mRNA levels (detected by qRT-PCR) at different time points during ESC–NPC differentiation. Mean ± SD; n=3 independent experiments. **c**, Quantification of the percentage of PAX6^+^ cells at different time points during ESC–NPC differentiation. Mean ± SD; n=3 independent experiments. **d**, Schematic illustration of the FUCCI system (upper panel) and images of FUCCI ESCs at indicated times of neural induction stained with DAPI and the PAX6 antibody (lower panels). FUCCI signals (GEMININ, green; CDT1, white) indicate cell cycle status. Scale bar, 10 μm.

**Extended Data Fig. 4.**
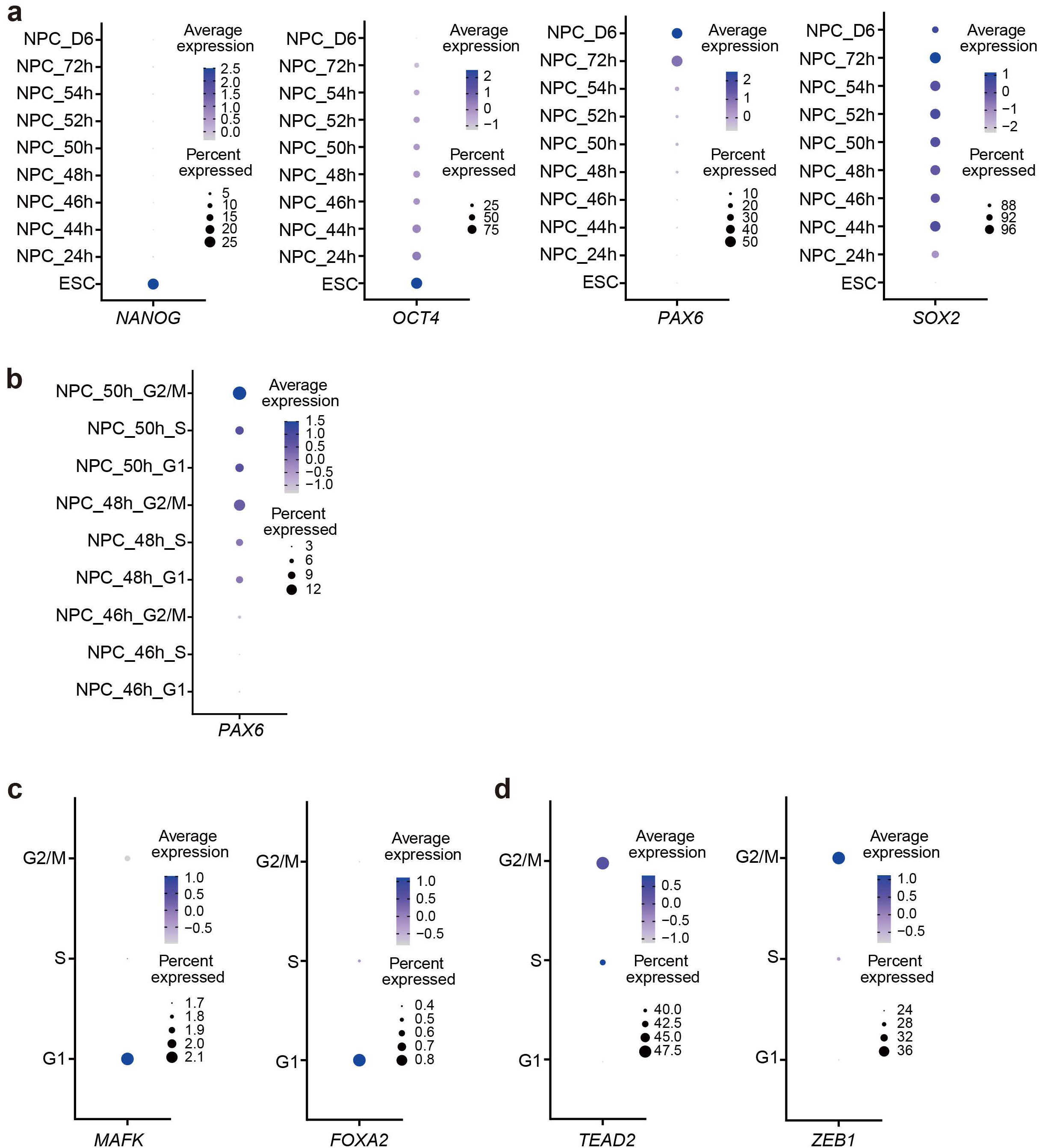
| Single-cell analysis of gene expression from NPC_D2 to NPC_D3. **a**, Dot blot showing the expression of *NANOG*, *OCT4*, *PAX6*, and *SOX2* from ESCs to NPC_D6 based on the single-cell RNA sequencing (scRNA-seq) data. **b**, Dot blot showing the cell cycle distribution of *PAX6*-positive cells from NPC_46h to NPC_50h. **c**,**d**, Dot blots showing two representative genes that are highly expressed in the G1 phase (**c**) or the G2 phase (**d**) based on the integrated scRNA-seq data.

**Extended Data Fig. 5.**
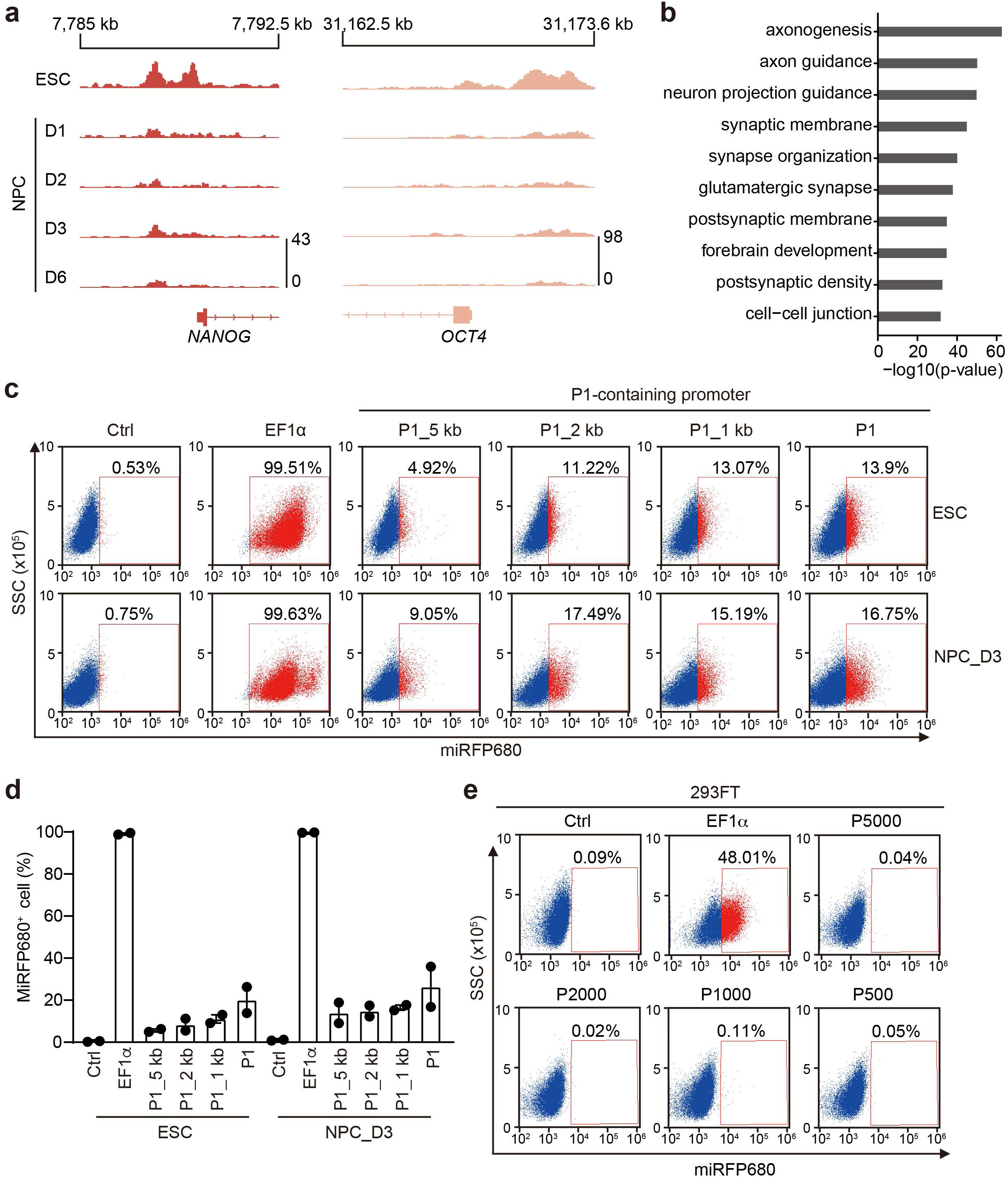
| G2-specific PAX6 transcription is driven by the 500-bp promoter. **a**, ATAC-seq tracks showing the chromatin accessibility at the *NANOG* and *OCT4* gene loci in ESCs and NPCs from days 1–6 of neural induction. **b**, Top 10 enriched GO pathways of genes with increased chromatin accessibility in NPC_D3 compared to NPC_D2. **c**, FACS plots showing the expression of the miRFP680 reporter driven by *PAX6* P1-containing promoters of different lengths in ESCs and NPCs at day 3 of neural induction (NPC_D3). The EF1α promoter was used as the positive control. Uninfected cells (Ctrl) were used as the negative control. **d**, Quantification of the percentage of miRFP680^+^ cells in **c**. Mean ± range; n=2 independent experiments. **e**, FACS plots showing the expression of the miRFP680 reporter driven by P500-containing promoters in 293FT cells. The EF1α promoter was used as the positive control, and uninfected cells (Ctrl) were used as the negative control.

**Extended Data Fig. 6.**
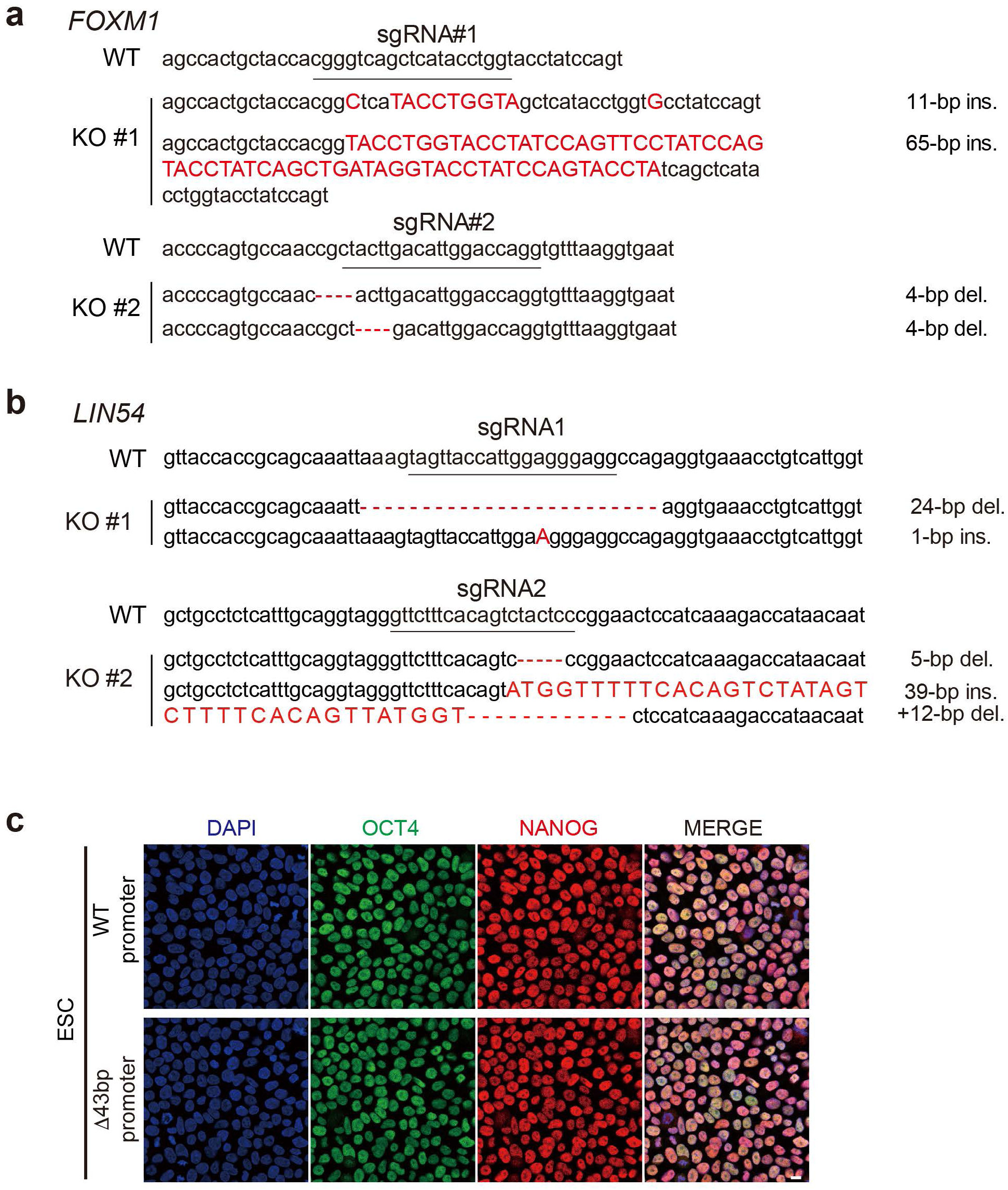
| Characterization of genome-edited ESC clones. **a**,**b**, Sanger sequencing of the *FOXM1* (**a**) and *LIN54* (**b**) genes in different ESC knockout clones. The sgRNA sequences used for the genome editing were underlined in the wild-type (WT) reference sequence. Indels and insertions in the genome were shown in red font. **c**, Images of WT ESCs or ESCs with the Δ43bp deletion from the *PAX6* promoter stained with DAPI and antibodies against OCT4 (green) and NANOG (red). Scale bar, 10 μm.

**Extended Data Fig. 7.**
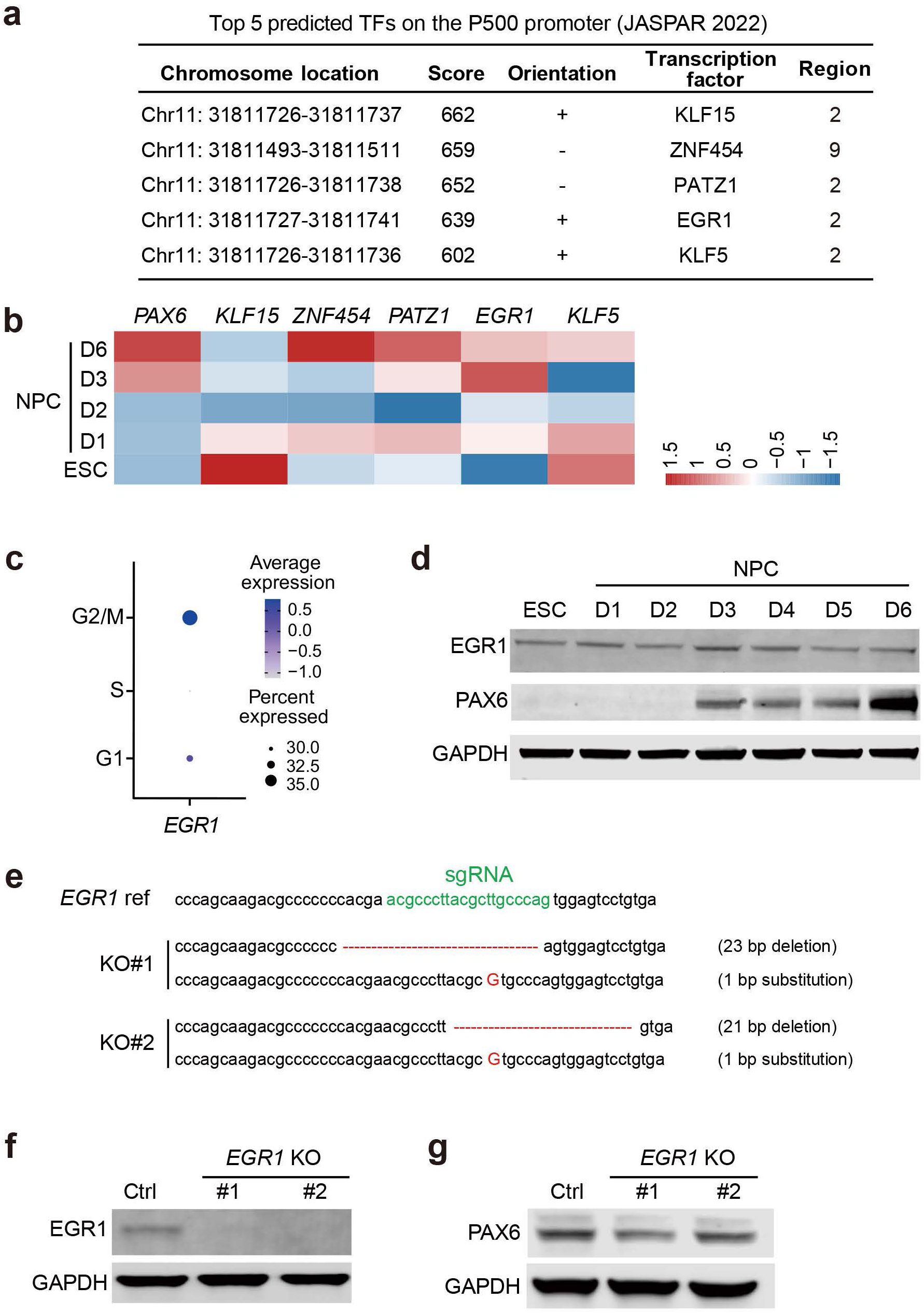
| EGR1 is not required for *PAX6* expression during ESC–NPC transition. **a**, Top 5 transcription factors (TFs) predicted to target the *PAX6* 500-bp promoter. KLF15, PAZT1, EGR1, and KLF5 are predicted to target the 2nd 50-bp segment, whereas ZNF454 is predicted to target the 9th 50-bp segment. **b**, Heatmap showing the expression values of *PAX6*, *KLF15*, *PATZ1*, *EGR1*, *KLF5*, and *ZNF454* in ESCs and NPCs at different times of neural induction. **c**, Dot plot showing the cell cycle distribution pattern of *EGR1* expression in ESCs and NPC_52h based on the integrated scRNA-seq data. **d**, Western blots showing the protein levels of EGR1 and PAX6 in ESCs and NPCs. **e**, Sanger sequencing of the *EGR1* gene in two ESC knock out clones. The sgRNA sequence was highlighted in green, and indels in the genome were indicated in red. **f,g**, Western blots showing the protein levels of EGR1 in WT and *EGR1* KO ESCs (**f**) and PAX6 in WT and *EGR1* KO NPCs at day 3 of neural induction (**g**).

**Extended Data Fig. 8.**
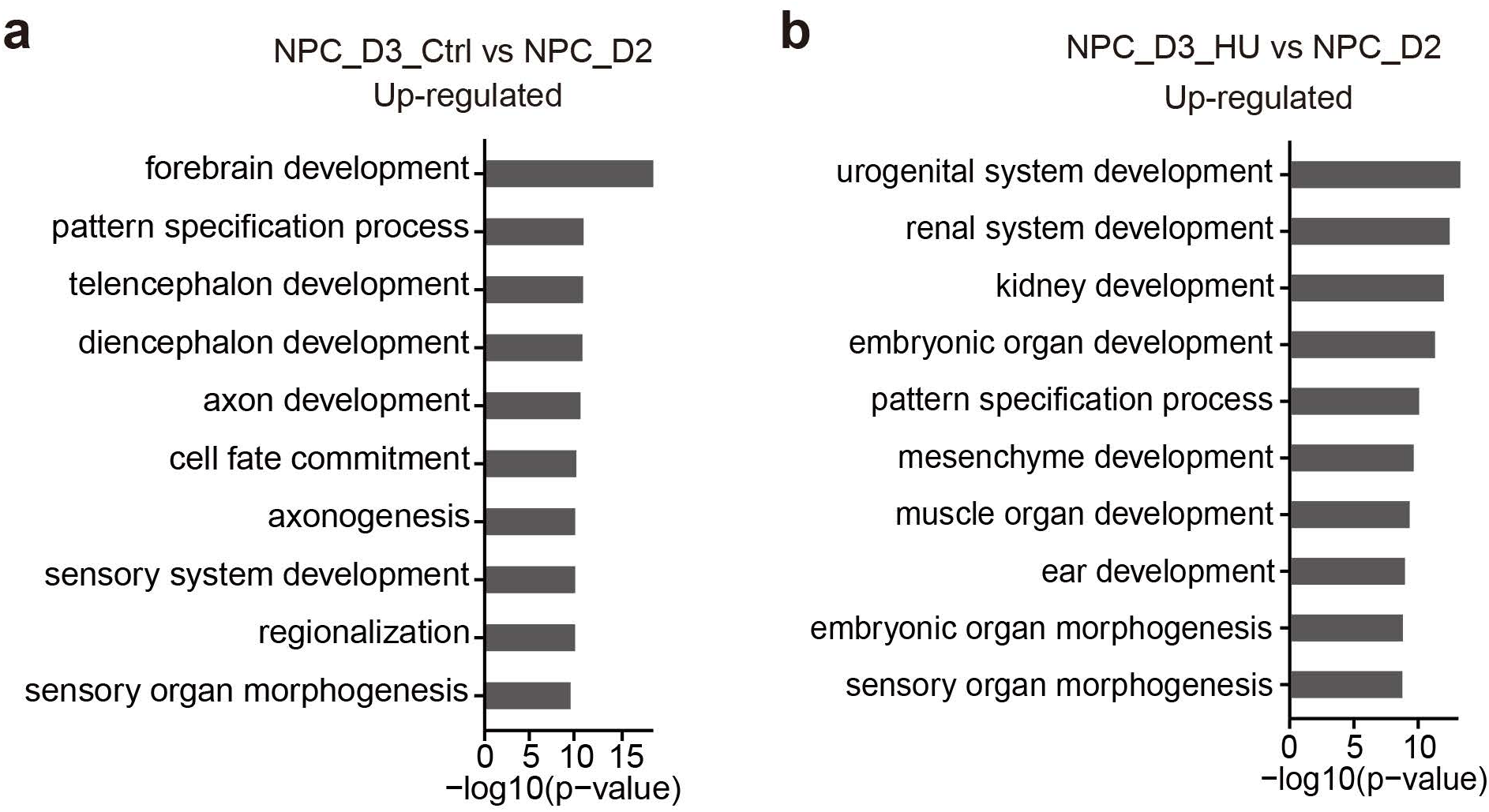
| Hydroxyurea (HU) treatment impairs the ESC–NPC transition. **a**, Top 10 enriched GO pathways of upregulated genes in normally differentiated NPC_D3 compared to NPC_D2. **b**, Top 10 enriched GO pathways of upregulated genes in HU-treated NPC_D3 compared to NPC_D2.

